# CV-containing vesicle budding from chloroplast inner envelope membrane is mediated by clathrin but inhibited by GAPC

**DOI:** 10.1101/2021.12.02.470903

**Authors:** Ting Pan, Yangxuan Liu, Chengcheng Ling, Yuying Tang, Wei Tang, Yongsheng Liu, Liang Guo, Chanhong Kim, Jun Fang, Honghui Lin, Eduardo Blumwald, Songhu Wang

## Abstract

Clathrin-mediated vesicular formation and trafficking are highly conserved in eukaryotic cells and are responsible for molecular cargo transport and signal transduction among organelles. It remains largely unknown whether clathrin-coated vesicles can be generated from chloroplasts. *CHLOROPLAST VESICULATION* (*CV*)-containing vesicles (CVVs) generate from chloroplasts and mediate chloroplast degradation under abiotic stress. In this study, we showed that CV interacted with the clathrin heavy chain (CHC) and induced vesicle budding from the chloroplast inner envelope membrane. Defects on *CHC2* and the dynamin-encoding *DRP1A* gene affected CVV budding and releasing from chloroplast. *CHC2* is also required for CV-induced chloroplast degradation and hypersensitivity to water stress. Moreover, GLYCERALDEHYDE-3-PHOSPHATE DEHYDROGENASE (GAPC) interacts with CV and impairs the CV-CHC2 interaction. *GAPC1* overexpression inhibited CV-mediated chloroplast degradation and hypersensitivity to water stress. *CV* silencing alleviated the hypersensitivity of *gapc1gapc2* plant to water stress. Together, our work revealed a pathway of clathrin-assisted CVV budding from the chloroplast inner envelope membrane, which mediated the stress-induced chloroplast degradation and stress response.

## Introduction

Endocytosis and vesicular trafficking are indispensable for eukaryotic cells. By mediating protein transport and protein degradation, they facilitate key cellular processes, such as signal transduction, cell polarization, responses to stimuli, etc (Fan, Li et al., 2015, Paez Valencia, Goodman et al., 2016). Clathrin-mediated endocytosis (CME) is the primary endocytic mechanism for the uptake of membrane proteins, lipids and extracellular molecules into the plant cell. The process of CME is highly conserved among species; it is initiated by the adaptor protein (AP) complexes and accessory proteins interacting with membrane lipids and cargo proteins, recruiting clathrin to the membrane (Fan et al., 2015, Paez Valencia et al., 2016). Clathrin is composed of three clathrin heavy chains (CHCs) and three clathrin light chains (CLCs), forming three-legged triskelions that self-polymerize (Kirchhausen, 2009). There are two functionally redundant *CHC* genes (*CHC1* and *CHC2*) (Kitakura, Vanneste et al., 2011) and three *CLC* genes (*CLC1-CLC3*) in the Arabidopsis genome (Wang, Yan et al., 2013a). Mutations in clathrin genes influence plant development (Kitakura et al., 2011, Wang et al., 2013a, Yu, Zhang et al., 2016), stomatal movements (Larson, Van Zelm et al., 2017) and plant immunity (Wu, Liu et al., 2015). For instance, the developmental abnormality of the mutants *chc2*, *clc2*, and *clc3* was caused by the impaired endocytosis of the PIN auxin transporter (Kitakura et al., 2011, Wang et al., 2013a, Yu et al., 2016). Following the clathrin coat assembly, the mature clathrin-coated vesicles are released into the cytosol through membrane scission, a process dependent on the enzyme dynamin. Dynamin-related proteins (DRP1 and DRP2) participate in clathrin-mediated trafficking, including CME (Fujimoto, Arimura et al., 2010, Konopka, Backues et al., 2008). The DRP family of proteins colocalize with CLCs at the cell plasma membrane and are required for cell polarization and cytokinesis in Arabidopsis (Fujimoto et al., 2010, Konopka et al., 2008).Glyceraldehyde-3-phosphate dehydrogenase (GAPDH) is a highly-conserved enzyme involved in glycolysis that catalytically converts glyceraldehyde-3-phosphate to 1,3-bisphosphoglyceric acid. The Arabidopsis genome contains several different types of GAPDH isoforms (GAPA, GAPB, GAPCp1, GAPCp2, and GAPC1 and GAPC2) which reside in different subcellular compartments. In spite of their fundamental roles in carbon metabolism, cytoplasmic GAPC1 and GAPC2 display moonlight functions (Zaffagnini, Fermani et al., 2013), including non-metabolic roles in regulating ABA-induced stomatal closure (Guo, Devaiah et al., 2012), mitochondria metabolism (Schneider, Knuesting et al., 2018, Zaffagnini et al., 2013), ROS (Henry, Fung et al., 2015), autophagy (Han, Wang et al., 2015, Henry et al., 2015), and even accumulating in the nucleus to regulate gene expression (Kim, Guo et al., 2020, Testard, Da Silva et al., 2016, Vescovi, Zaffagnini et al., 2013, Zhang, Zhao et al., 2017).

Endocytosis and vesicle trafficking facilitate material exchange and signaling between the different cell compartments (Fan et al., 2015, Paez Valencia et al., 2016). However, the connection(s) between chloroplasts and other organelles remains to be elucidated. During natural or stress-induced leaf senescence, chloroplastic Nitrogen-rich proteins undergo vacuolar-mediated degradation, enabling N-remobilization to sink tissues (Masclaux, Valadier et al., 2000). To date, multiple pathways of chloroplast degradation have been characterized (Otegui, 2018, Xie, Michaeli et al., 2015). The first reported pathway is mediated by senescence-associated vacuoles (SAVs) which are lytic compartments containing proteases such as SENESCENCE ASSOCIATED 12 (SAG12) (Carrion, Costa et al., 2013, Martinez, Costa et al., 2008, Otegui, Noh et al., 2005). Next, three autophagy-dependent pathways, including Rubisco-containing bodies (RCBs) (Ishida & Yoshimoto, 2008, Ishida, Yoshimoto et al., 2008, Wada, Ishida et al., 2009), ATG8-interacting Protein 1 (ATI1-PS) bodies (Michaeli & Galili, 2014, Michaeli, Honig et al., 2014) and small starch-like granule (SSLG) bodies (Wang & Liu, 2013, Wang, Yu et al., 2013b), were shown to be involved in the turnover of chloroplasts and starch. These autophagic bodies are transported to the vacuole for eventual degradation (Xie et al., 2015).

Previously, we characterized a gene *CHLOROPLAST VESICULATION* (*CV*) encoding a protein involved in a pathway that was independent of autophagy and SAVs (Wang & Blumwald, 2014). Natural- and stress-induced senescence induced *CV* expression and CV targeted chloroplasts, mediating the formation of CV-containing vesicles, termed CVVs, that carried chloroplast proteins into the vacuole for degradation (Wang & Blumwald, 2014). CV silencing increased plant tolerance to water stress through the enhancement of nitrogen assimilation in rice (Sade, Umnajkitikorn et al., 2018). Although the formation of CVVs was observed and their role in mobilizing chloroplast proteins was partially characterized, the mechanism(s) mediating the CVV formation remained largely unknown. Here, we show that CV interacts clathrin to generate vesicles from chloroplast inner membrane and that the membrane scission of the CVVs is mediated by DRP1A. In addition, GAPC interacts with CV to disrupt CVV formation and thereby inhibit CV-induced chloroplast degradation under water stress.

## Results

### CV interacts with clathrin heavy chain

To elucidate mechanisms related to CV-induced vesicle formation, we identified proteins associated with CV via co-immunoprecipitation (co-IP) and liquid chromatography-tandem mass spectrometry (LC-MS/MS). CV-expression was driven by the dexamethasone (DEX)-inducible promoter DEX in transgenic *DEX-CV-HA* plants (Wang & Blumwald, 2014). Proteins were extracted from transgenic *DEX-CV-HA* plants after treatment with DEX and Concanamycin A, an inhibitor of endocytic vesicle trafficking (Dettmer, Hong-Hermesdorf et al., 2006) that limited the transport of CVVs out of the chloroplast (Wang & Blumwald, 2014). CV-HA was immunoprecipitated using agarose beads conjugated with monoclonal antibodies raised against the HA tag. Clathrin heavy chain 2 (CHC2), a key component of CME, was repeatedly identified in the three independent experiments (Table SI).

To confirm the CV–CHC2 interaction, we conducted bimolecular fluorescence complementation (BiFC) (Figure 1). Transient expression of both fusion genes *CV-nYFP* and *cYFP-CHC2* in tobacco mesophyll protoplasts resulted in BiFC fluorescence that was observed in the dots nearby the chloroplasts (Figure 1A). Co-expression of *nYFP-CV* and *cYFP-CHC2* also showed positive BiFC signals, although these signals were distributed in the cytosol (Figure 1B), possibly because the N-terminal tag nYFP disrupted the CV-chloroplast transit signal peptide, misdirecting CV to the cytosol. Similar patterns were observed when the constructs of *GFP-CV* and *CV-GFP* were expressed separately in protoplasts (Figure S1A). *GFP-CV* was localized to the cytosol without an obvious dot-like structure (Figure S1A), similar to the observation of BiFC signal between *nYFP-CV* and *cYFP-CHC2* (Figure 1B). Furthermore, BiFC also indicated that the interaction between CV and CHC1 (Figure 1C), which shares a 98% amino acid similarity with CHC2, also occurred in the similar dot-like structures.

**Figure 1:**
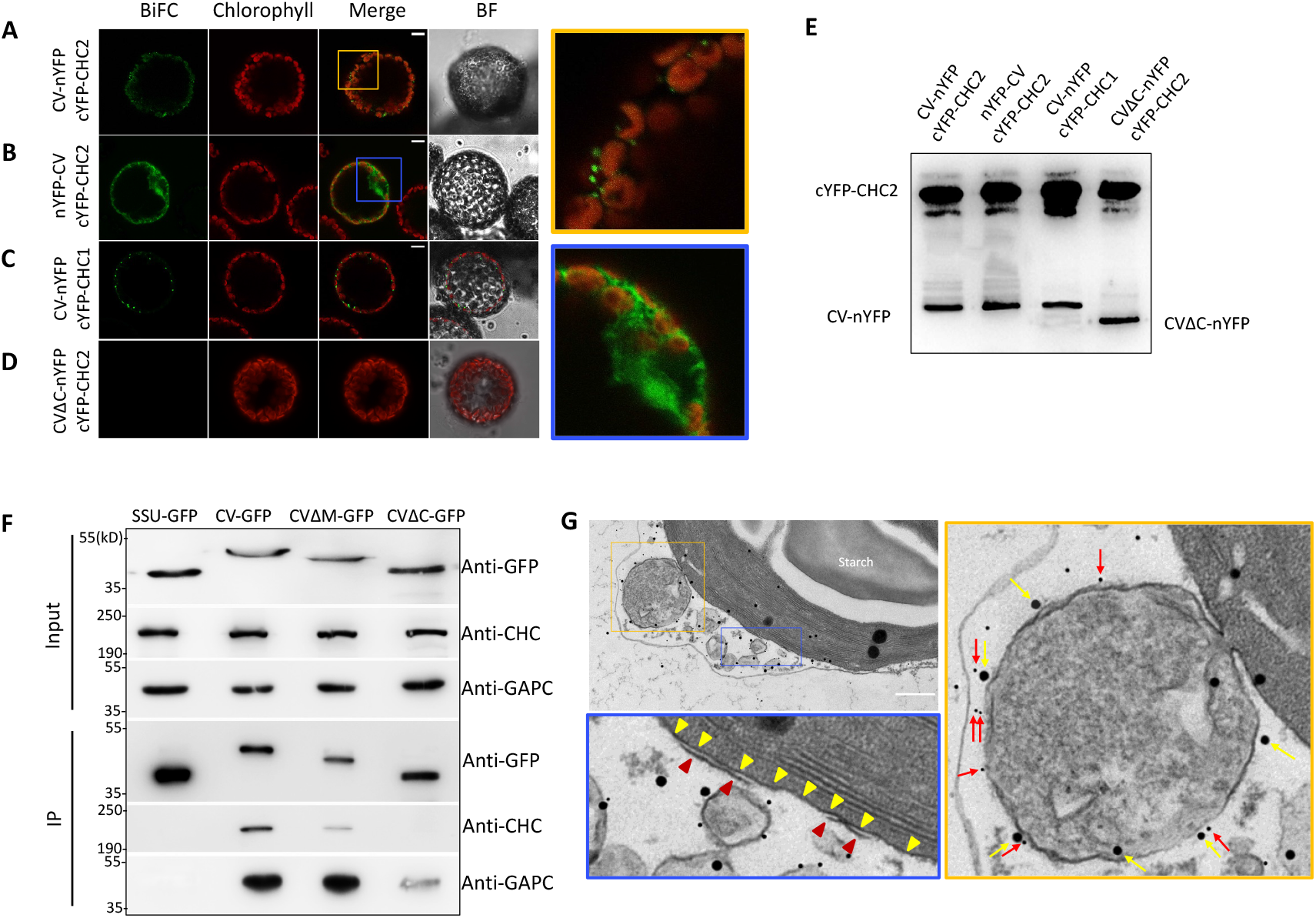
CV interact with CHC in vivo. (A) to (D) BiFC analysis of in vivo interaction: CV-nYFP and cYFP-CHC2 (A), nYFP-CV and cYFP-CHC2 (B), CV-nYFP and cYFP-CHC1 (C), and CVΔC-nYFP and cYFP-CHC2 (D). BiFC, yellow fluorescence from interacting nYFP and cYFP; Chlorophyll, autofluorescence of chlorophyll; BF, bright field. Bar=10μm. Right panels are expanded from the orange and blue insets in (A) and (B), respectively. (E) Total proteins were extracted from the transformed protoplasts (A-D) and immunoblot assays were perfomed by the anti-GFP polyclonal antibodies. (F) Co-IP assays with GFP-tagged CV, CVΔM, CVΔC, and SSU. Crude lysates (Input) from transgenic plants expressing SSU-GFP, CV-GFP, CVΔM-GFP, and CVΔC-GFP were immunoprecipitated with an anti-GFP antibody. Co-IP samples were detected by immunoblotting with the anti-GFP, anti-CHC, and anti-GAPC antibodies, respectively. (G) Immunodetection of CV-HA (yellow arrows) and CHC (red arrows) in the vesicles budding from chloroplasts. The section was obtained from 2-week-old seedlings of CPro::CV-GFP plants treated with 100mM mannitol and 3µM Concanamycin A for 12h. The panels with orange or blue frame are zoom-in observations of the top left panel. In the orange-frame panel, yellow and red arrows indicate CV-HA and CHC, respectively. In the blue-frame panel, yellow and red triangles indicate the inner and outer envelope membranes, respectively. Bar=300nm.

In addition to the N-terminal transit signal peptide, which determines its chloroplast localization, CV protein contains another two domains: a transmembrane domain and a highly conserved C-terminus domain (Figure S1B). In order to determine the domain responsible for the interaction of CV with CHC, we used transgenic plants expressing *DEX-CV-GFP*, *DEX-CVΔTM* (deletion of the transmembrane domain)-*GFP*, and *DEX-CVΔC* (deletion of the C-terminus conserved domain)-*GFP* (Wang & Blumwald, 2014) to perform co-IP assays. Transgenic plants expressing the gene encoding the chloroplast-localized small subunit of ribulose-1,5-bisphosphate carboxylase (SSU) fused to *GFP* was used as a control (Kim, Meskauskiene et al., 2012). The co-IP assays showed that the deletion of C-terminus conserved domain completely abolished the interaction between CV and CHC, as indicated by immunoblotting with anti-CHC antibodies that recognize both CHC1 and CHC2 (Figure 1F). Consistently, no BiFC signal was observed in protoplasts co-expressing *CVΔC* and *CHC2* (Figure 1D), although immunoblot showed that both genes are substantially expressed (Figure 1E). To further visualize colocalization of CV and CHC *in vivo*, we used native promoter of *CV* gene (CPro)(Wang & Blumwald, 2014) to drive *CV-GFP* fusion gene and Col-0 plants were transformed by the construct CPro::CV-GFP. CV-GFP was detected by immunoblot (Figure 3A) or confocal observation (Figure 3B and 3C) in the *CPro::CV-GFP* transgenic plants under 200mM mannitol-caused osmotic stress but not under control condition, which is consistent with our previous study showing that *CV* is induced by abiotic stress (Wang & Blumwald, 2014). Under native promoter, CV-GFP dots are also associated with or released from chloroplasts (Figure 3C), which is consistent with the observation of transient expression of *35S::CV-GFP* in protoplast (Figure S1A) and with the previous results of DEX-treated *DEX-CV-GFP* transgenic plants (Wang & Blumwald, 2014). Next, we used the leaf sections from *CPro::CV-GFP* plants treated with moderate osmotic stress and Concanamycin A to perform the double immunolabeling with anti-GFP and anti-CHC antibodies. Transmission electron microscopy (TEM) observations indicated the association of CHC dots (red arrows in orange frame of Figure 1G) with the CV-labeled vesicles (yellow arrows in Figure 1G) budding from chloroplast membrane. CV and CHC antibodies also labeled some vesicles that were not associated with chloroplasts and perhaps released from chloroplasts (Figure 1G). Time-lapse observation of protoplasts transiently expressing *CV-GFP* and *mCherry-CHC2* showed that initially, CV-GFP formed “loci” first (purple triangles at “0” second, Figure S2) and that mCherry-CHC2 gradually concentrated on the loci (white and blue arrows, Figure S2). These results indicated that CV and CHC2 are co-localized in the vesicles budding from chloroplast.

**Figure 2:**
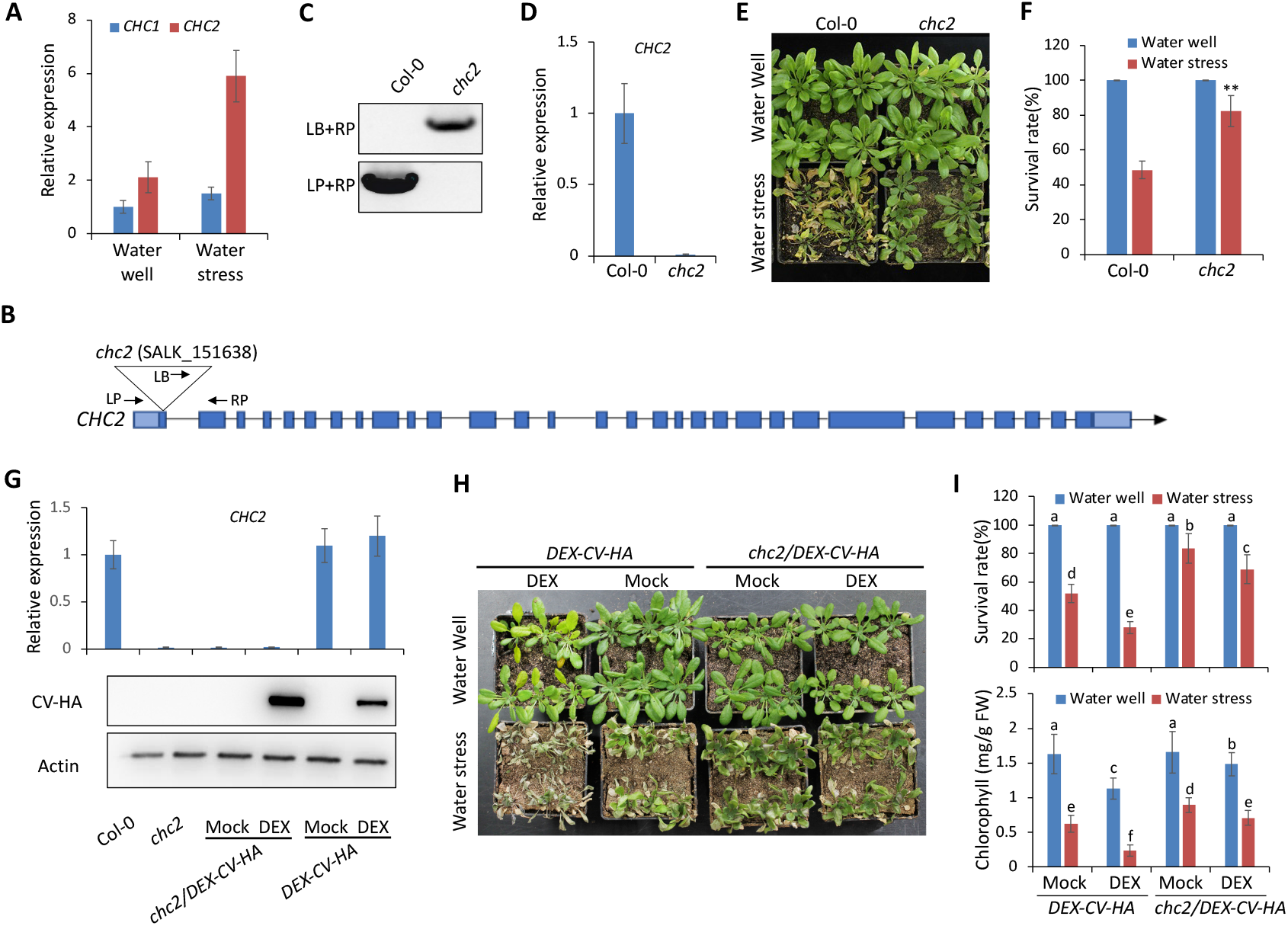
T-DNA mutation of *CHC2* suppressed CV’s role in chloroplast degradation and water stress response. (**A**) qRT-PCR assays of *CHC1* and *CHC2* expression in 10-day-old Col-0 plants with or without water stress. (**B**) The T-DNA of mutant *chc2* (germplast SALK_151638) was inserted in the coding region of first exon of *CHC2* gene. Dark and light blue boxes indicated the coding regions and untranslated regions, respectively. (**C** and **D**) Genotyping of T-DNA insertion (**C**) and qRT-PCR analysis of *CHC2* expression (**D**) in mutant *chc2*. (**E** and **F**) The phenotypes (**E**) and survival rates (**F**) of Col-0 and *chc2* mutant subjected to water stress by withholding water for 14 days and re-watered for 3d. Sixteen plants for each line were evaluated and Means ± SD were obtained from three independent tests. Asterisks “**” indicate significant difference at P<0.001. (**G**) qRT-PCR analysis of *CHC2* expression and immunoblot with anti-HA and anti-Actin antibodies in Col-0, *chc2*, *chc2/DEX-CV-HA*, and *DEX-CV-HA* plants. The *chc2/DEX-CV-HA* and *DEX-CV-HA* plants were treated without (Mock) or with 10μM DEX for 24h. (**H** and **I**) The phenotypes (**H**), survival rates and chlorophyll contents (**I**) of 2-week-old *chc2/DEX-CV-HA* and *DEX-CV-HA* plants subjected to water stress by withholding water for 12 days. Sixteen plants for each line were evaluated and Means ± SD were obtained from three independent tests. Asterisks “**” indicate significant difference at P<0.001.

**Figure 3:**
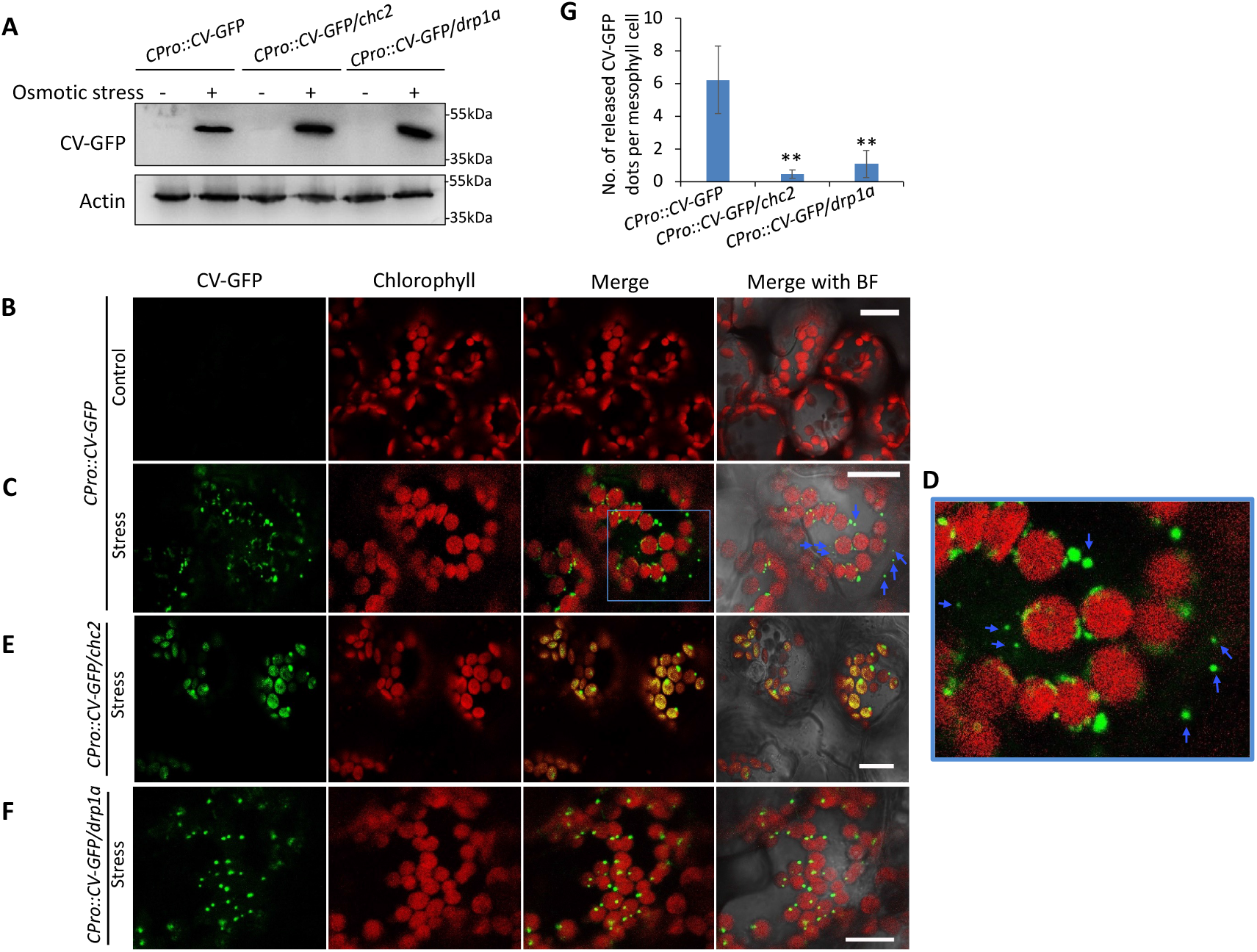
CVV formation and release from chloroplast is disrupted by defects on *CHC2* and *DRP1A* gene. (**A**) Immunoblot analysis of 2-week-old transgenic seedlings of *CPro::CV-GFP*, *CPro::CV-GFP/chc2*, and *CPro::CV-GFP/drp1a* treated without or with 200mM mannitol for 24h. Total proteins were extracted and immunodetected by anti-GFP and anti-Actin antibodies. (**B-D**) Confocal microscope observation of mesophyll cells from 2-week-old transgenic seedlings of *CPro::CV-GFP* treated without (**B**) or with (**C**) 200mM mannitol for 24h; (**D**) Zoom-in image from the blue frame in (**C**); GFP was showed in the green and chlorophyll autofluorescence was showed in the red. BF means bright field. Blue arrows indicate the CV-GFP-containing vesicles (CVVs) released from chloroplasts. Bar=10μm for all the images. (**E** and **F**) Confocal microscope observation of mesophyll cells from 2-week-old transgenic seedlings of *CPro::CV-GFP/chc2* (**E**) and *CPro::CV-GFP/drp1a* (**F**) treated with 200mM mannitol for 24h. (**G**) The average number of CV-GFP dots released from chloroplast per mesophyll cell of *CPro::CV- GFP, CPro::CV-GFP/chc2*, and *CPro::CV-GFP/drp1a* plants under osmotic stress. More than 50 mesophyll cells for each line were observed and means ± SD were obtained from three independent tests. Asterisks “**” indicate significant difference at P<0.001.

### *CHC2* is required for CV-induced chloroplast degradation and hypersensitivity to water stress

Our quantitative reverse transcription-PCR (qRT-PCR) assays indicated that mRNA levels of *CHC2* is higher than that of *CHC1* in Col-0 plants, especially under water stress (Figure 2A). Therefore, we isolated the homozygous T-DNA insertion line *chc2* (SALK_151638) (Figure 2B and 2C). The T-DNA is inserted in the coding sequence in the first exon of *CHC2* (Figure 2B) and no obvious expression of *CHC2* was detected (Figure S2C) by qRT-PCR, suggesting that *chc2* is a knockout mutant. Under water stress, the *chc2* plants showed the increased survival rate than Col-0 plants (Figure 2E and 2F), which is consistent with a previous study (Larson et al., 2017). To assess the genetic CV-CHC2 interaction, we crossed *chc2* mutant with *DEX-CV-HA* line. The *chc2/DEX-CV-HA* line was validated by qRT-PCR and immunoblot assays (Figure 2G). DEX treatment induced chloroplast degradation in *DEX-CV-HA* plants, as indicated by the yellowing leaves (Figure 2H) and decreased chlorophyll contents (Figure 2I) under water well conditions. However, DEX-induced senescence was impaired in *chc2/DEX-CV-HA* line (Figure 2H and 2I). Under water stress, DEX treatment decreased chlorophyll contents and survival rates in *DEX-CV-HA* plants, while *chc2/DEX-CV-HA* plants displayed higher chlorophyll contents and higher survival rates than *DEX-CV-HA* plants, with or without DEX treatment (Figure 2H and 2I). These results indicated that *CHC2* is required for CV-induced chloroplast degradation and hypersensitivity to water stress.

Next, the *chc2* plants were transformed by CPro::CV-GFP vector to produce the transgenic line *CPro::CV-GFP/chc2*, respectively. CV-GFP was detected in *CPro::CV-GFP/chc2* line under 200mM mannitol-caused osmotic stress but not under control condition, similar to *CPro::CV-GFP* line (Figure 3A). In *CPro::CV-GFP* mesophyll cells, CV-GFP signals were observed to be associated with chloroplasts and some CV-GFP-labeled dots, as indicated by the blue arrows in Figures 3C and 3D, were apart from chloroplasts. Those dots released from the chloroplast are named CV-containing Vesicles (CVVs). We observed more than 50 mesophyll cells of *CPro::CV-GFP* plants from three independent experiments to calculate the number of released CVVs per cell. The results showed that there were about 6 released CVVs in every mesophyll cell (Figure 3G). In *CPro::CV-GFP/chc2* cells, however, most CV-GFP signals were overlapped with chlorophyll autofluorescence (red) (Figure 3E). A few released CVVs were also observed in some mesophyll cells but the mean value was much lower than that in *CPro::CV-GFP* plants (Figure 3G). Immunolabeling TEM analysis showed that the CV-GFP-labeled vesicles were budding from chloroplast inner envelope membrane in *CPro::CV-GFP* plants (Figure 1G). However, no budding vesicle was observed in the *CPro::CV-GFP/chc2* plants under stress (Figure 4A) but some CV-concentrated loci (yellow frame in Figure 4A), which might be the above-mentioned dot-like structure shown in the confocal microscopy (Figure 3E). CV-GFP also labeled thylakoid membranes (blue frame in Figure 4A), which is consistent with the confocal observation that most CV-GFP signals are overlapped with chlorophyll (Figure 3E). These results indicate that *CHC2* plays an important role in the CV-mediated vesicle budding and formation.

**Figure 4:**
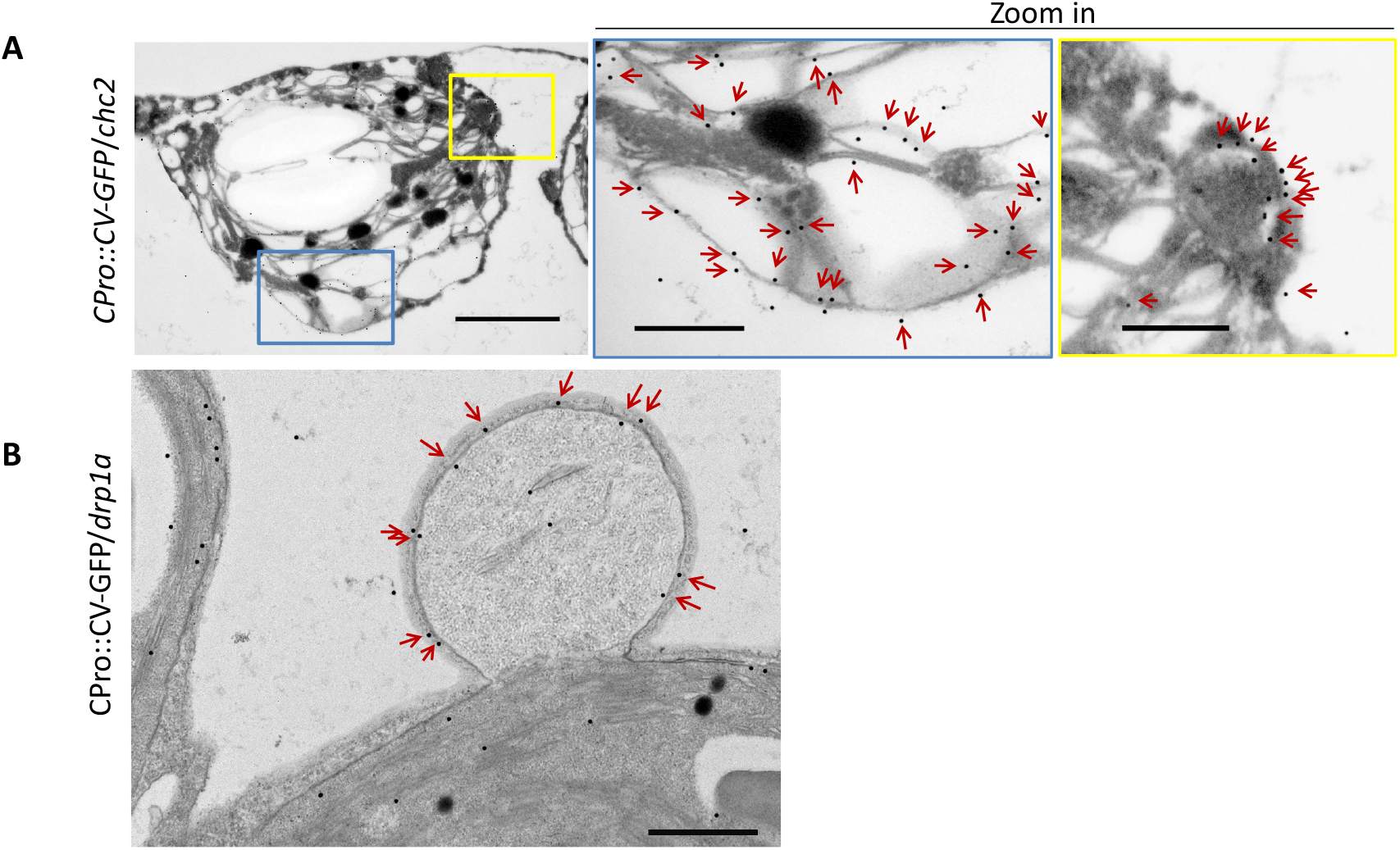
Immunolabeling TEM analysis of CV-GFP in leaf sections of *CPro::CV-GFP/chc2* and *CPro::CV-GFP/drp1a*. (**A** and **B**) Immunolabeling TEM analysis of CV-GFP in the leaf sections of 20-day-old seedlings of *CPro::CV-GFP* transformed *chc2* (**A**) and *drp1a* (**B**) plants treated with 200mM mannitol for 24h. The ultrathin sections from these leaves were immunolabeled by anti-GFP antibodies before TEM observation. The middle and right panels in (**A**) were amplified from blue and yellow frame in the left panel, respectively. The black gold particles, as indicated by the red arrows, showed the CV-GFP localization. Bars in the left, middle, right panel of (**A**) indicates 1μm, 200nm, and 200nm, respectively. Bar in the (**B**) indicates 200nm.

### Defect on *DRP1A* genes interfered with CVV release from chloroplasts

Dynamin participates in the formation and release of clathrin-coated vesicles (Holstein, 2002). We assessed the subcellular localization of CV and of dynamin-related protein 1A (DRP1A). *CV-GFP* and *mCherry-DRP1A* were transiently expressed in Col-0 mesophyll protoplasts and fluorescence was assessed by confocal microscopy. Some of the red fluorescence of mCherry-DRP1A overlapped with CV-GFP dots, while the rest of the red signal was localized in the cytosol (Figure S3A). By contrast, no obvious overlapping between CV-GFP and mCherry was detected (Figure S3A). Partial colocalization of CV-GFP and mCherry-DRP1A were further quantified by calculating Pearson (Rp) and Spearman’s (Rs) correlation coefficients (Figure S3A) via software ImageJ with PSC colocalization plugin (French, Mills et al., 2008). These results implied a possible role of DRP1A in the membrane scission and release of CVVs. We isolated *drp1a,* the homologous line of T-DNA insertion mutant of *DRP1A* gene (SALK_069077) (Figures. S3B and S3C). Although the T-DNA was inserted in an intron of *DRP1A* gene, its expression was almost silenced as indicated by qRT-PCR showing that *DRP1A* mRNA levels in the *drp1a* mutant were about 5% of that of WT plants (Figure S3D). The *drp1a* mutant was much smaller than the Col-0 (Figure S3E), suggesting that the mutation of *DRP1a* has a great impact on plant growth and development. The transgenic *CPro::CV-GFP/drp1a* line was generated and immunoblotting analysis indicated that CV-GFP was also induced by the osmotic stress in the *CPro::CV-GFP/drp1a* line (Figure 3A). Confocal observation showed that CV-GFP formed dot-like structures (Figure 3F), which might be the vesicles budding from chloroplast and the TEM observation confirm the existence of CV-GFP-label vesicle budding from chloroplast envelope membrane (Figure 4B). Very few CV-GFP dots released from chloroplast were also observed in some mesophyll cells of *CPro::CV-GFP/drp1a* line but the average level is much lower than that in *CPro::CV-GFP* plants (Figure 3G). These results suggested that *DRP1A* is required for CVV release from chloroplasts.

### CV is localized in the inner envelope membrane of chloroplast

We have shown previously that CV associated with thylakoid and chloroplast envelope membranes (Wang & Blumwald, 2014). In order to assess whether CV associated with the inner or outer chloroplast envelope membranes, we isolated chloroplasts from leaves of *DEX-CV-GFP* transgenic plants treated with DEX, separated the inner and outer membranes and assessed CV presence. Antibodies raised against GLUTAMINE SYNTHASE (GS), TIC110, and TOC75 were used as markers of chloroplast stroma, inner, and outer envelope membrane, respectively (Figure 5A). Anti-GS antibodies recognized both the chloroplastic (GS2) and cytosolic (GS1) isoforms, and no GS1 contamination was detected in the membrane fractions, indicating no cytosolic contamination. GFP-CV was mainly associated with the inner envelope membrane, although there was a slight trace at the outer membrane fraction (Figure 5A).

**Figure 5:**
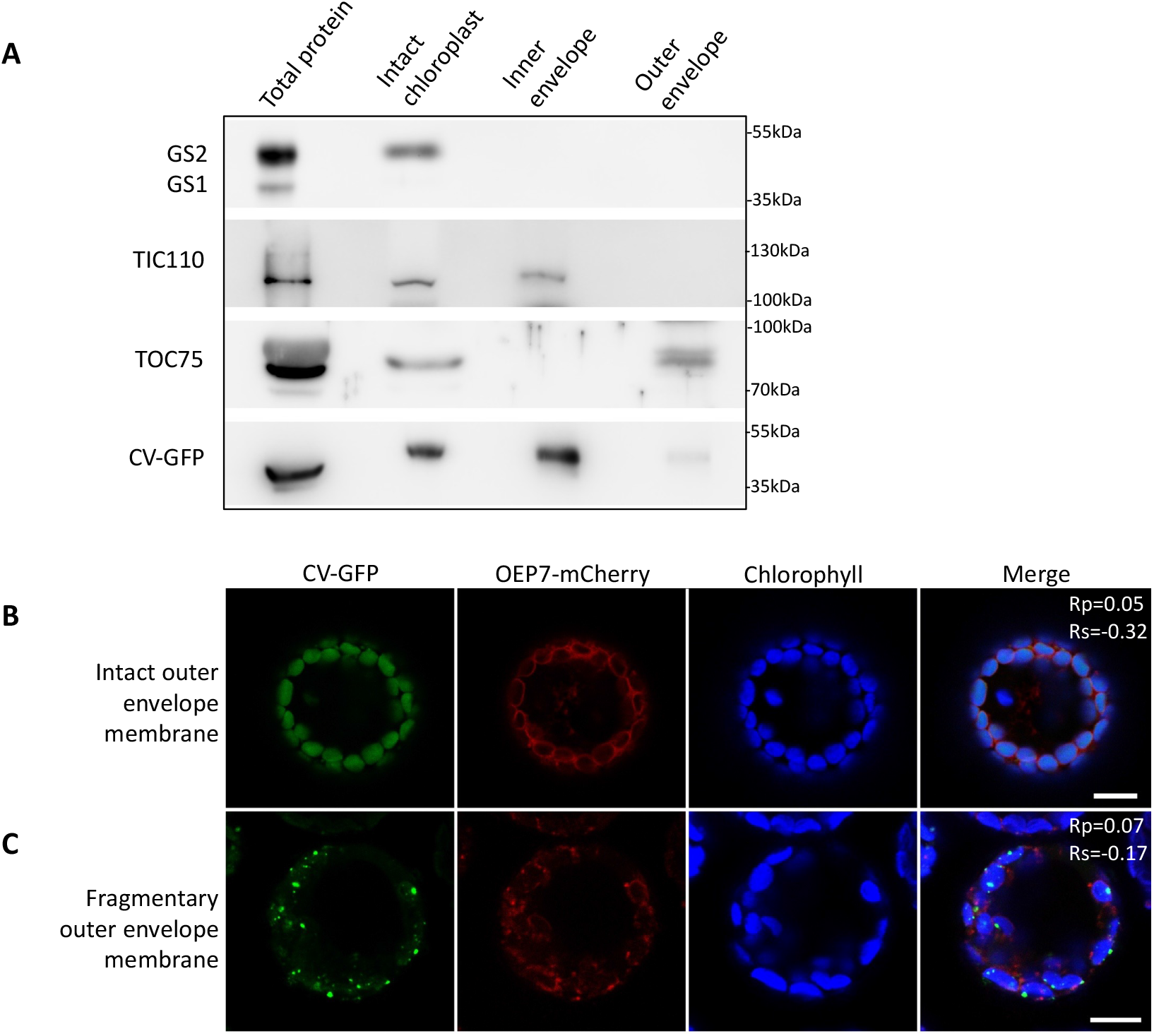
CV is localized in the inner envelope membrane of chloroplast. (**A**) Total proteins, intact chloroplasts, inner chloroplast envelope membranes, and outer chloroplast envelope membranes were isolated from leaves of DEX-treated *DEX-CV-GFP* transgenic plants and immunoblotted with antibodies raised against glutamine synthase (GS), TIC110, TOC75, and GFP, respectively. Anti-GS antibodies recognized both the chloroplastic (GS2) and cytosolic (GS1) isoforms. (**B** and **C**) The protoplasts isolated from wild type leaves were co-transformed by vectors of CV-GFP and *OUTER ENVELOPE PROTEIN 7* (*OEP7*) fused to mCherry. The outer chloroplast envelope membranes were intact in some transformed protoplasts (**B**) but fragmentary in the others (**C**), as labeled by OEP7-mCherry. Colocalization analysis of “green” and “red” channels was performed and Pearson’s (Rp) and Spearman’s (Rs) coefficients were calculated (n>10) and showed in the merge channel. The Rp and Rs values which were lower than 0.4 or negative indicate lack of colocalization.

Given the inner envelope membrane-localization of CV and the cytosolic localization of CHC2 (Song, Lee et al., 2006), the formation of CVVs would be limited by the outer chloroplast membrane. To assess the conditions for CVV formation, we transiently co-transformed protoplasts with *OUTER ENVELOPE PROTEIN 7* fused to *mCherry* (*OEP7-mCherry*) and *CV-GFP* (Figure 5B and 5C). CV-GFP fluorescence (green) overlapped with the chlorophyll autofluorescence (blue) and no obvious dot-like structure was observed in transformed protoplasts with an intact chloroplast outer envelope membrane (Figure 5B). In chloroplasts displaying a fragmented outer membrane (as indicated by OEP7-mCherry), CV-GFP-labeled dots were observed around the chloroplast (Figure 5C). Moreover, CV-GFP did not localize at the outer membrane since the green fluorescence of CV-GFP and the red OEP7-mCherry fluorescence did not overlap, which was confirmed by the very low Rp and Rs values (Figure 5C). The TEM microscopy also indicated that the CV-labeled vesicle was budding from the inner envelope membrane (as indicated by the yellow triangles in the blue frame of Figure 1G) when outer membrane was fragmented (as indicated by the red triangles in Figure 1G). These results supported the notion that fragmentation of the chloroplast outer membrane facilitated the CV-mediated vesicle budding possibly by making CHC2 accessible. Confocal microscopy indicated that more CV-GFP-labeled dot-like structures were observed in leaves from DEX-induced *DEX-CV-GFP* plants after mannitol treatment (Figure S4A). Co-IP assays confirmed that osmotic stress enhanced the CV-CHC2 interaction (Figure 6D and S5B).

**Figure 6:**
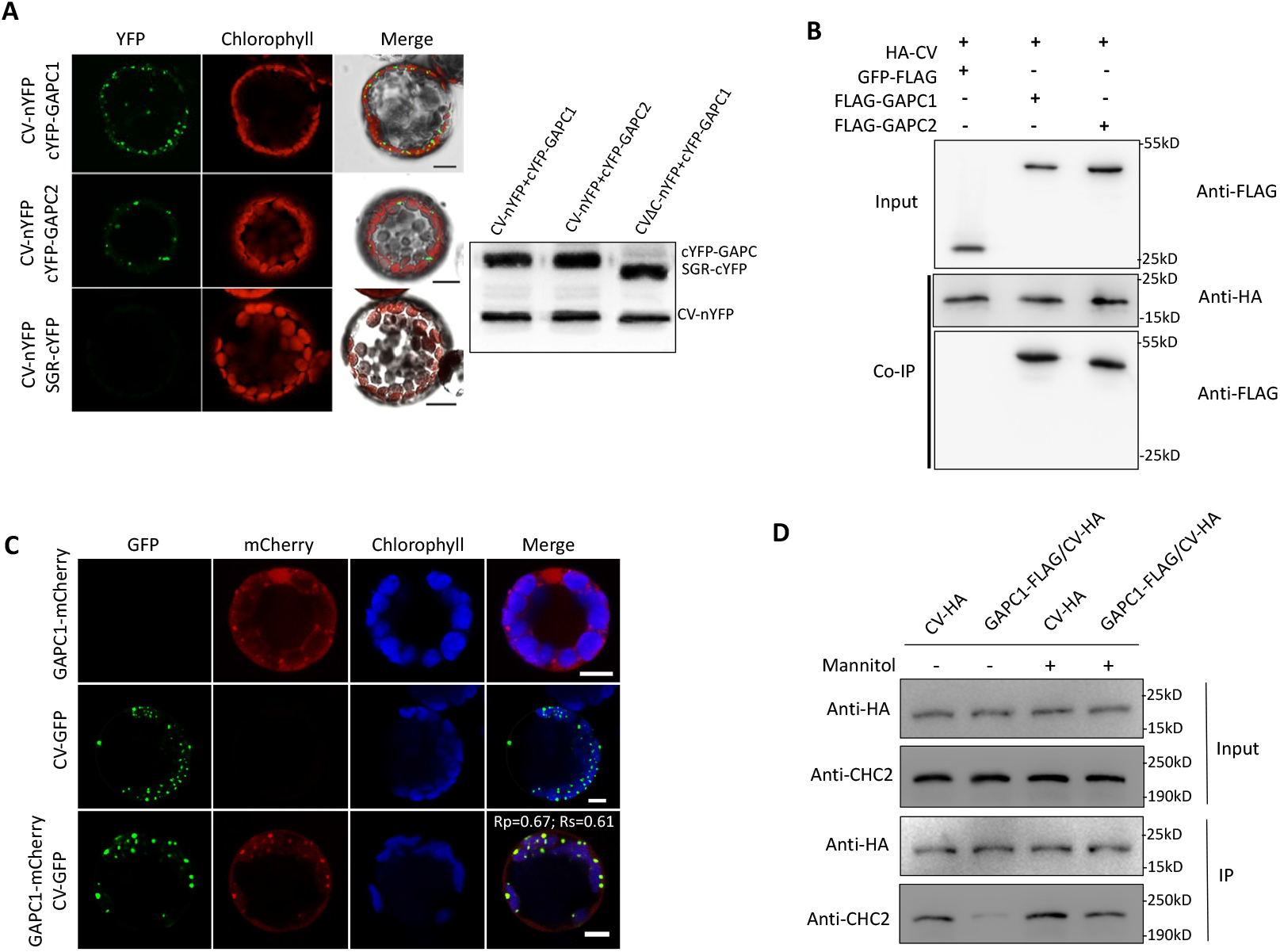
GAPC interacts with CV to inhibit CV-CHC2 interaction. (**A**) BiFC analysis of *in vivo* CV-GAPC interaction: CV-nYFP and cYFP-GAPC1, CV-nYFP and cYFP-GAPC2, and CV-nYFP and cYFP-SGR. BiFC, yellow fluorescence from interacting nYFP and cYFP; Chlorophyll, autofluorescence of chlorophyll. Bar=10μm. Total proteins were extracted from the transformed protoplasts and immunoblot assays was performed by using anti-GFP polyclonal antibodies. (**B**) Co-IP assays with HA-tagged CV. Crude lysates (Input) from tobacco leaves transiently expressing CV-HA plus GFP-FLAG, GAPC1-FLAG, and GAPC2-FLAG, respectively, were immunoprecipitated with anti-HA antibody. Co-IP samples were detected by immunoblotting with anti-HA and anti-FLAG antibodies, respectively. (**C**) Confocal microscopic observation of the wild type protoplasts transiently expressing CV-GFP or GAPC1-mCheery, or both. Bar=5μm. Colocalization analysis of CV-GFP and GAPC1-mCherry was performed. Pearson’s (Rp) and Spearman’s (Rs) coefficients were calculated (n>10) and showed in the “merge” channel of the bottom panel. The Rp and Rs values in the range +0.4 to 1 indicate colocalization. (**D**) Co-IP assays of CV-HA. Crude lysates (Input) from DEX-induced transgenic plants DEX-CV-HA and GAPC1-FLAG/DEX-CV-HA plants treated with or without 200mM mannitol for 24h, were immunoprecipitated with anti-HA antibodies. Co-IP samples were detected by immunoblotting with anti-HA and anti-CHC antibodies, respectively.

### GAPC interacts with CV to inhibit CV-CHC2 interaction

Co-immunoprecipitation of CV and LC-MS/MS-based identification of proteins interacting with CV showed that GAPC1 was a putative interactor of CV (Table SI). Recent studies indicated that GAPC1, in addition to its glycolytic properties, is an enzyme with many moonlighting functions (Wojtera-Kwiczor, Gross et al., 2013, Zaffagnini et al., 2013, Zhang et al., 2017), including a positive role in water stress (Guo et al., 2012). BiFC (Figure 6A) and co-IP assays (Figure 6B) confirmed the interactions between CV and two functionally redundant GAPC isoforms (GAPC1 and GAPC2), sharing 98% amino acid sequence identity (Guo et al., 2012). We used antibodies raised against GAPC, which were validated by immunoblotting analysis of the *GAPC* overexpression line and double mutants (Figure S5), to evaluate the CV-GAPC interaction (Figure 1F). Our results indicated that the CV transmembrane domain was not required for the interaction, but the deletion of the CV conserved domain obviously impaired the interaction (Figure 1F). The transient expression of GAPC-mCherry and CV-GFP also confirmed that GAPC-mCherry was not only localized at the cytosol, but also partially associated with CV-GFP (Figure 6C). Since both GAPC and CHC2 interact with CV, we further analyzed whether GAPC overexpression affects CV-CHC2 interaction. Co-IP assays of DEX-treated *DEX-CV-HA* and *GAPC1-FLAG/DEX-CV-HA* plants (Figure 7A) with or without 200mM mannitol showed that osmotic stress enhanced CV-CHC2 interaction and *GAPC1* overexpression inhibited CV-CHC2 interaction in the absence or presence of osmotic stress (Figure 6D). These results indicated that *GAPC* overexpression impairs CV-CHC2 interaction.

**Figure 7:**
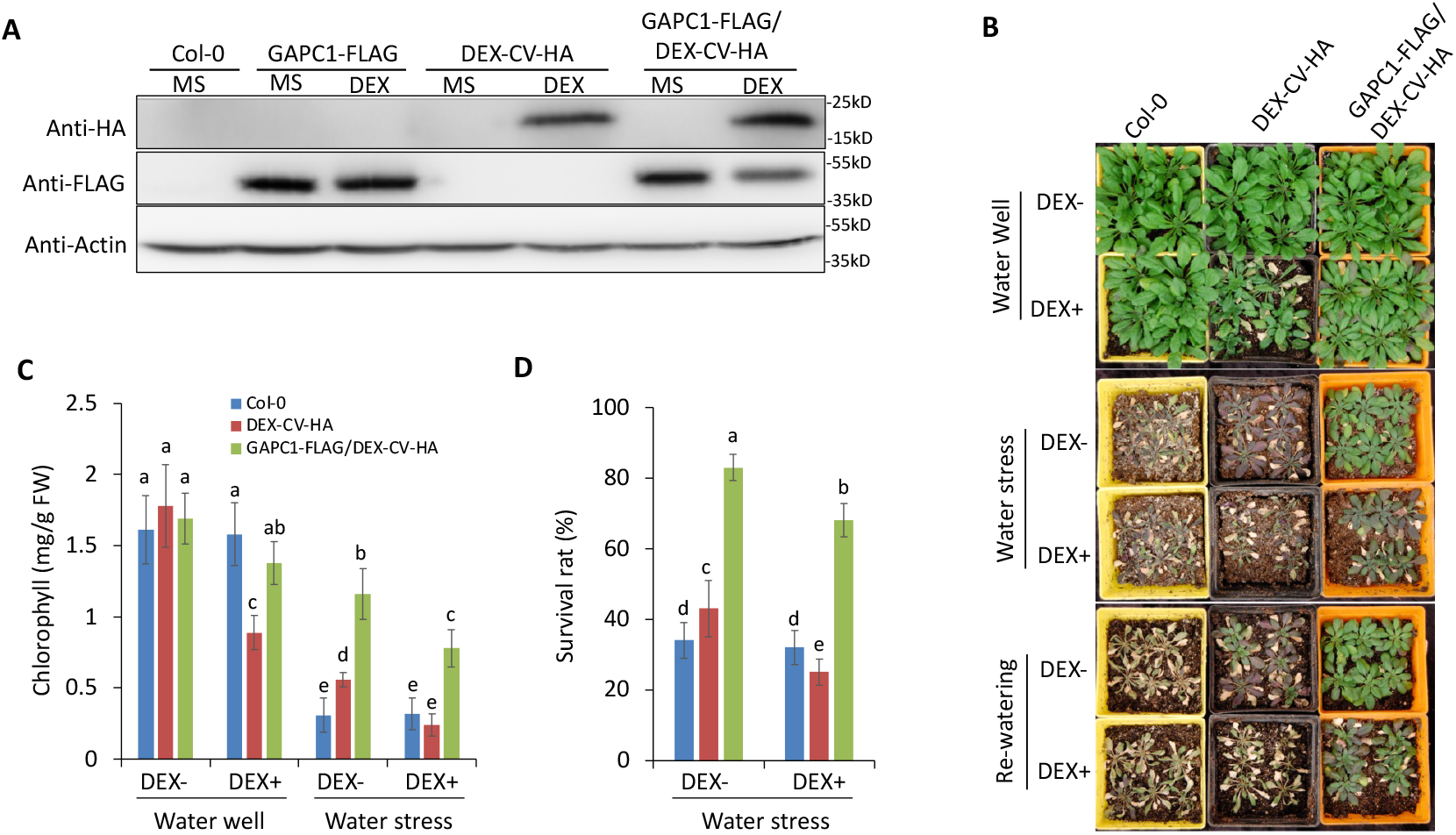
GAPC overexpression inhibits CV-induced chloroplast degradation and hypersensitivity to water stress. (**A**) Total proteins extracted from Col-0, *GAPC1-FLAG, DEX-CV-HA* transgenic lines, and the cross line *GAPC1-FLAG/DEX-CV-HA* were immunoblotted with anti-HA, anti-FLAG, and anti-Actin antibodies. (**B**) Two-week-old seedlings of Col-0, *GAPC1-FLAG-, DEX-CV-HA-*transgenic lines and the cross line *GAPC1-FLAG/DEX-CV-HA* were subjected to water stress for 14d by withholding water and re-watered for 3d. (**C** and **D**) Total chlorophyll content (**C**) and Survival rate (**D**) were determined under well-watered conditions and 3 d after re-watering, as described in (**B**). Eighteen plants for each line were evaluated and means ± SD were obtained from three independent experiments. Different letters indicate significant differences (Duncan test, P<0.05).

### Genetic interaction between CV and GAPC1

To investigate the genetic interaction between *CV* and *GAPC*, *GAPC1-FLAG/DEX-CV-HA* double-overexpression (DO) lines were generated (Figure 7A) by crossing *DEX-CV-HA-1* and *GAPC1-FLAG-1* plants (Figure S6). DEX treatment induced chloroplast degradation in DEX-CV-HA plants, as indicated by the senescent leaves (Figure 7B) and decreased chlorophyll contents (Figure 7C) under water well conditions. However, DEX-induced senescence was suppressed in the DO line (Figure 7B and 7C). Under water stress, DEX treatment decreased chlorophyll contents and survival rates in *DEX-CV-HA* plants, while *GAPC1-FLAG/DEX-CV-HA* plants displayed higher chlorophyll contents and higher survival rates than *DEX-CV-HA* plants, with or without DEX treatment (Figure 7C and 7D). These results indicated that GAPC inhibited CV-induced chloroplast degradation and hypersensitivity to water stress. Moreover, we generated the triple knockdown line *amiR-CV-1/gapc1/gapc2,* by crossing the *CV*-silenced line *amiR-CV-1* (Wang & Blumwald, 2014) with the double *GAPC* knockout *gapc1-1/gapc2-1* (Figures 8A and 87B). While *gapc1-1/gapc2-1* plants displayed water stress hypersensitivity (Guo et al., 2012) (Figs 8C and 8D), the *amiR-CV-1/gapc1/gapc2* plants showed higher survival rates (Figure 8D) and chlorophyll contents (Figure 8E), suggesting that *CV* silencing alleviated the hypersensitivity of *gapc1-1/gapc2-1* plants to water stress.

**Figure 8:**
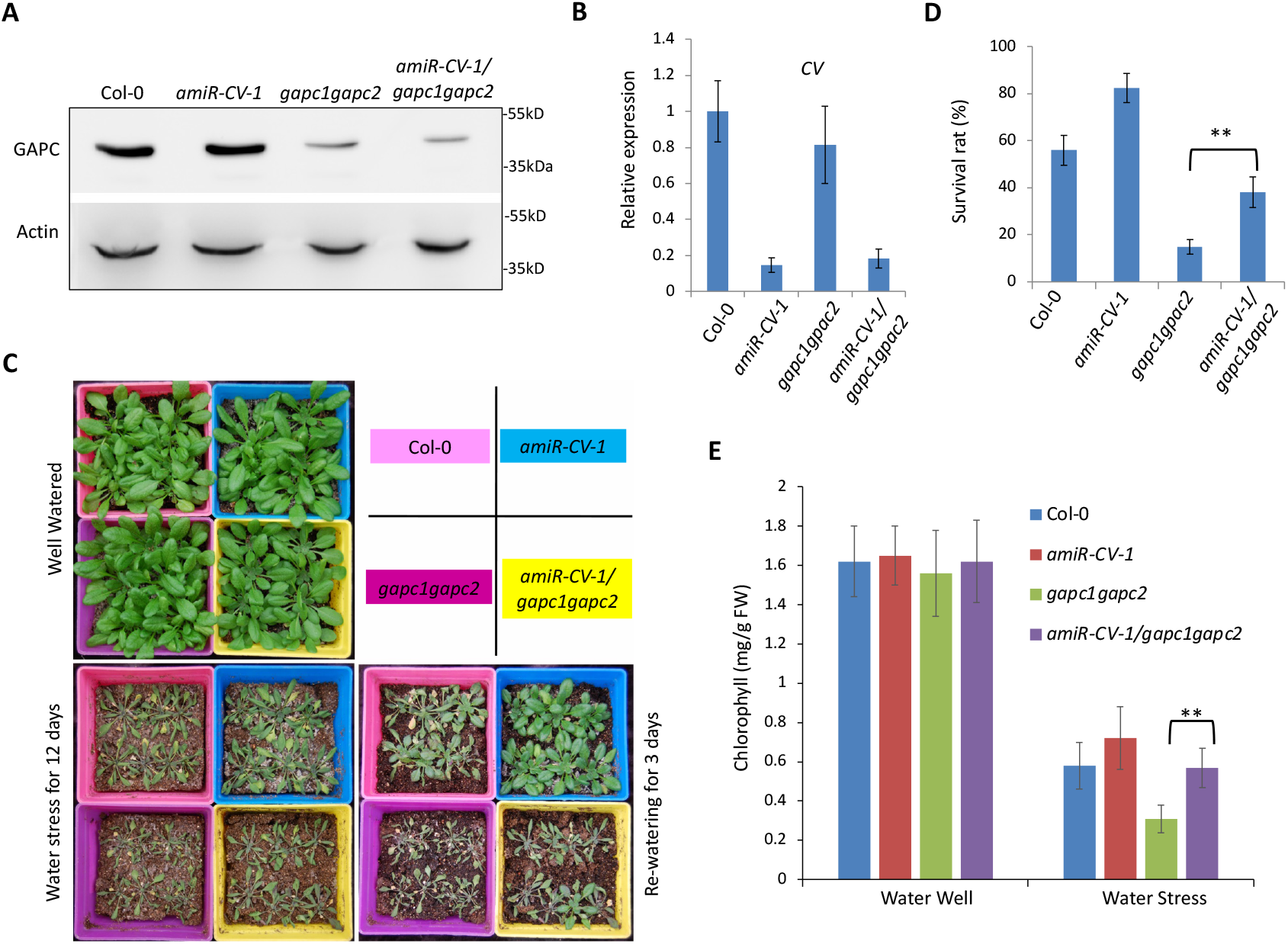
CV silencing alleviates the drought hypersensitivity of *gapc1gapc2* plants. (**A**) Total proteins extracted from 3-week-old seedlings of Col-0, *amiR-CV-1*, *gapc1gapc2*, and the crossed line *amiR-CV-1/gapc1gapc2* were immunoblotted with anti-GAPC and anti-Actin antibodies. (**B**) The qRT-PCR analysis of CV expression in 3-week-old seedlings of Col-0, *amiR-CV-1*, *gapc1gapc2*, and *amiR-CV-1/gapc1gapc2*. (**C**) Two-week-old seedlings of Col-0, *amiR-CV-1*, *gapc1gapc2*, and *amiR-CV-1/gapc1gapc2* were subjected to water stress for 12 d followed by rewatering for 3 d. (**D** and **E**) Survival rate (**D**) and chlorophyll contents (**E**) were determined under well-watered conditions and 3 d after rewatering, as described in (**C**). Sixteen plants for each line were evaluated and means ± SD were obtained from three independent tests. Asterisks “**” indicate significant difference at P<0.001.

## Discussion

Endocytosis and vesicular trafficking are highly conserved in eukaryotic cells and are responsible for molecular cargo transport and signal transduction between organelles. To date, the characterization of the mechanisms mediating the connection between the chloroplast and other intracellular compartments is scanty. We have shown previously that a protein encoded by *CV,* could mediate the vesicle formation from chloroplasts and these CVVs were involved in the degradation of stromal proteins at the plant vacuole and were associated with stress-induced chloroplast degradation (Sade et al., 2018, Wang & Blumwald, 2014). However, the mechanism(s) regulating the role of CV in vesicle formation from the chloroplast remains unknown. Similar to animal cells, plant cells contain three main types of coated vesicles: clathrin-coated vesicles (CCVs), coat protein I (COPI)-coated vesicles, and COPII-coated vesicles (Hwang & Robinson, 2009). Currently, it is well known that CCVs in plant cells are generated at two locations (Robinson & Pimpl, 2014) - the plasma membrane (PM) and the *trans*-Golgi network (TGN) - with the assistance of different adaptor protein (AP) complexes. Clathrin-mediated endocytosis from PM is dependent on AP2 and TPLATE complexes (Bashline, Li et al., 2013, Di Rubbo, Irani et al., 2013, Fan, Hao et al., 2013, Gadeyne, Sanchez-Rodriguez et al., 2014, Kim, Xu et al., 2013, Yamaoka, Shimono et al., 2013). CCVs budding from TGN is mediated by AP1 (De Marcos Lousa, Gershlick et al., 2012, Park, Song et al., 2013), AP3 complexes (Feraru, Paciorek et al., 2010, Lee, Kim et al., 2007, Zwiewka, Feraru et al., 2011), and some non-classical adaptors (Sauer, Delgadillo et al., 2013, Song et al., 2006) and might be responsible for the export of soluble vacuolar cargo from TGN.

Although bioinformatic analysis showed that clathrin- and COPI-dependent vesicle systems were not present in the chloroplast (Lindquist, Alezzawi et al., 2014), our results indicated that CV interacted with clathrin to induce vesicle budding outwards from the chloroplast envelope membrane (Figure 9). CV interacted with CHC *in vivo* and the loss of the C-terminus conserved domain of CV completely abolished its interaction with CHC (Figure 1). Genetic evidence indicated that *CHC2* is required for CV-induced chloroplast degradation and hypersensitivity to water stress (Figure 2H and 2I). CV can interact with both CHC1 and CHC2 (Figure 1). Defect on *CHC2* has an obvious inhibition on the CV’s function, possibly because *CHC2* has higher expression levels than *CHC1* under stress condition (Figure 2A). Defects on *CHC2* and *DRP1A* genes impaired the CVV formation and departure from chloroplasts (Figure 3). Together, these results suggested that the CME pathway might be involved in CV-induced vesicle budding and release from the chloroplast. With respect to the CV-CHC interaction, we can speculate that CV might function as a non-classical adaptor protein (AP). However, the APs involved in CME are soluble proteins lacking a transmembrane domain. CV is structurally similar to a receptor-like cargo protein with a transmembrane domain (Paez Valencia et al., 2016). Further investigations are required for elucidating whether CV is a non-classical adaptor protein or a receptor-like cargo protein for chloroplast degradation.

**Figure 9:**
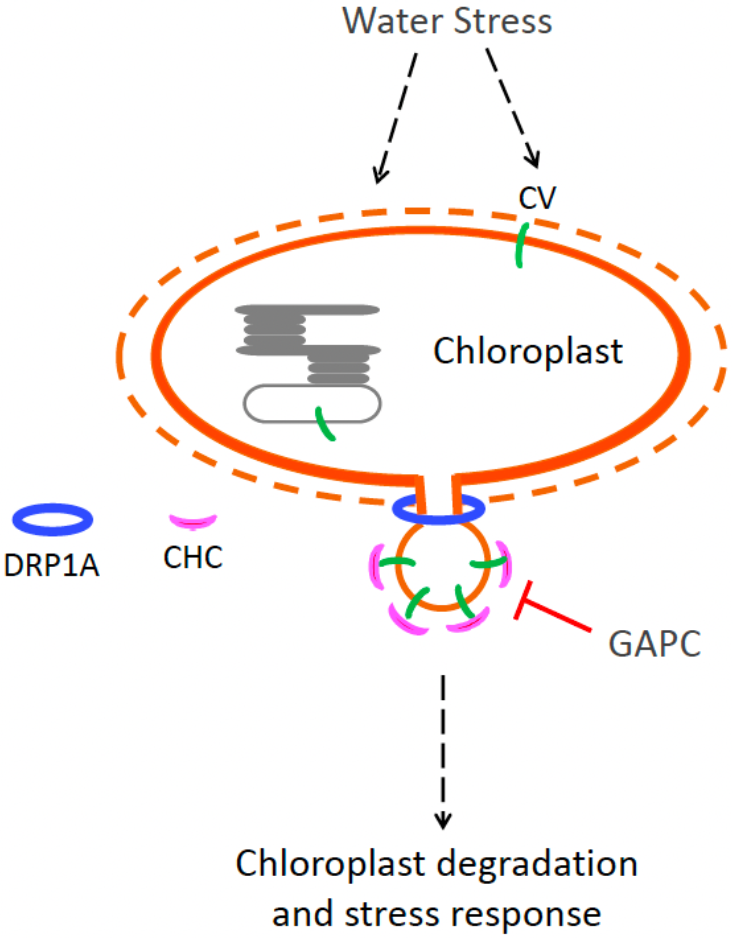
CVV budding from chloroplast envelope membrane was assisted by clathrin but inhibited by GAPC. Water stress induces *CV* gene expression and CV (Green) targets the chloroplast inner envelope membrane (Orange line) and thylakoid membrane (gray line). Envelope membrane-localized CV has access to cytosol-localized CHC (Pink) and possibly other clathrin structures for vesicle budding, because the chloroplast outer membrane (Orange dashed line) might be damaged or permeable under stress condition. The CV-containing vesicles (CVVs) are released from the chloroplast membranes after DRP1A (Blue) assembles around the neck of the budding vesicle, causing membrane scission. GAPC interacts with CV to impair CV-CHC interaction and thereby inhibits CV-mediated chloroplast degradation in plant response to water stress.

CV is associated with the chloroplast inner membrane (Figure 5A), consistent with the overlapping of CV-GFP and TIC20-II RFP, an inner membrane protein (Wang & Blumwald, 2014). This conclusion was further supported by the lack of overlapping of CV-GFP green fluorescence with the red signal of the outer envelope protein OEP7-mCherry (Figure 5C). Given the localization of CV at the inner chloroplastic membrane, the question of how CV can interact with the cytosol-localized CHC2 arises. Our analysis indicated that in the presence of an intact chloroplast outer membrane (Figure 5B) or in mutant *chc2* (Figure 3E), CV-GFP was uniformly distributed in the chloroplast and no CV-GFP-labeled dot-like structure was observed. However, damage or fragmentation of the outer envelope membrane facilitated CV-mediated vesicle budding formation (Figure 1G and 5C), possibly by increasing the accessibility to CHC and other clathrin-associated apparatus. Numerous studies showed the dynamic remodeling of the chloroplast outer membrane (Breuers, Brautigam et al., 2012, Breuers, Brautigam et al., 2011, Inoue, 2011) and its damage by abiotic stress (Fourrier, Bedard et al., 2008, Kwon, Verma et al., 2013, Moellering, Muthan et al., 2010, Wang, Uddin et al., 2014) or during leaf senescence (Springer, Kang et al., 2016). Under these conditions, *CV* expression is triggered (Wang & Blumwald, 2014) and CV can access clathrin and dynamin to mediate vesicle formation from the chloroplast inner envelope membrane. Our results confirmed that osmotic stress enhanced CV-CHC2 interaction (Figure 6D and S4B) and CVV formation (Figure S4A). CV also targets to the thylakoid membranes in *chc2* mutant (Figure 3E) or when the outer membrane is integrated (Figure 5B), as indicated by the overlapping of CV-GFP and chlorophyll. Immunolabeling TEM confirmed that CV-GFP are substantially associated with thylakoid membrane in *chc2* mutant (Figure 4A). A few CV-GFP protein associating with thylakoid membrane were also observed in the Col-0 and *drp1a* mutant (Figure 1G and 4B) and in our previous study (Wang & Blumwald, 2014). Further investigations are required for studying the functions of CV on the thylakoid membranes, which probably include disrupting the PSII complex by interacting with PsbO (Wang & Blumwald, 2014).

CV also interacts with GAPC1 and GAPC2 *in vivo* (Figure 6A and 6B) and GAPC overexpression impairs CV-CHC2 interaction (Figure 6D). It is possible that GAPC hinders the CV-CHC interaction through occupying the conserved domain that is also required for the CV-CHC interaction (Figure 1D). Another possibility is that the oxidative modification of GAPC can cause misfolding and protein aggregation (Zaffagnini, Marchand et al., 2019), which might disrupt the CV-CHC2 interaction. Further studies are required for testing the mechanism(s) by which GAPC inhibits CV’s function. GAPC is a catalytic enzyme with multiple moonlighting functions, including a positive role in plant response to water stress (Guo et al., 2012) and heat stress (Kim et al., 2020). *GAPC* overexpression inhibited CV-induced chloroplast degradation and hypersensitivity to water stress (Figure 7) and *CV* silencing alleviated the hypersensitivity of *gapc1gapc2* double mutant to water stress (Figure 8), suggesting that GAPC facilitate the plant response to water stress, in part, through suppressing CV’s function.

In addition to stress response, cytoplasmic GAPC1 and GAPC2 are also involved in mitochondria metabolism (Schneider et al., 2018, Zaffagnini et al., 2013), ROS (Henry et al., 2015), autophagy (Han et al., 2015, Henry et al., 2015), and gene expression (Testard et al., 2016, Vescovi et al., 2013, Zhang et al., 2017). In tobacco, GAPC interacted with autophagy-related protein 3 (ATG3) and silencing *GAPCs* activated ATG3-dependent autophagy (Han et al., 2015). Given the role(s) of autophagy in chloroplast degradation, GAPC might be a key regulator of chloroplast degradation. Based on our results, a model for the CV-mediated vesicle budding from the inner chloroplast envelope can be postulated (Figure 9). Water stress activates *CV* gene expression and also damages the outer chloroplast membrane. CV targets the chloroplast inner envelope membrane, as well as thylakoid membrane, and accesses the cytosol-localized CHC2 and other clathrin structures for vesicle budding. DRP1A is responsible for the CVV release from chloroplast. GAPC, a positive regulator of the stress response, interacts with CV to impair CV-CHC2 interaction and thereby inhibit CV-induced chloroplast degradation and hypersensitivity to water stress.

Together, our results revealed that the CV-containing vesicle budding from chloroplast inner envelope membrane is mediated by clathrin and GAPC interacts with CV to suppress CV-induced chloroplast degradation under water stress.

## Materials and Methods

### Plant Materials and Growth Conditions

*Arabidopsis thaliana* plants were grown in a chamber at 23°C under 16h-light/8h-dark cycles. Seeds were surface-sterilized and sowed on MS/2 medium (PhytoTechnology laboratories) (1% sucrose pH 5.7) and chilled at 4°C for 2 d. The ecotype Columbia-0 (Col-0) was used as wild type plants. The transgenic lines expressing an artificial microRNA targeting *CV* (*amiR-CV*), the line with a chemically-induced expression of *CV-GFP* (*DEX-CV-GFP*) and a mutagenesis deletion to remove either the chloroplast trans-membrane domain (*DEX-CVΔTM-GFP*) or the C-terminus conserved domain (*DEX-CVΔCD-GFP*) of CV, the line with a chemically-induced expression of *CV-HA* (*DEX-CV-HA*), and the line of *ProCV::GUS* was obtained as described previously(Wang & Blumwald, 2014). The double mutants (*gapc1-1gapc2-1*) were generated as described previously(Guo et al., 2012). The T-DNA insertion mutant *chc2* (SALK_151638) *drp1a* (SALK_069077) was obtained from TAIR (www.arabidopsis.org) and the homozygous lines were isolated and confirmed by genomic PCR and qRT-PCR. The native promoter of CV (CPro)(Wang & Blumwald, 2014) was fused with *CV-GFP* gene and CPro::CV-GFP was inserted into pEG100 vector. pEG100-CPro::CV-GFP construct was used to transform Col-0, *chc2*, and *drp1a* mutants, respectively.

To generate *35S::GAPC1-FLAG* lines, the cDNA of *GAPC1* was fused with *6×FLAG* by fusion PCR, and a Gly-Ala-linker was introduced between *GAPC1* and *6×FLAG*. The fusion fragment (*GAPC-linker-FLAG*) were cloned into the vector pBI121 by double digests reaction. The *pBI121-GAPC1-FLAG* construct was used to transform Col-0 wild type plants by the floral dipping method(Clough & Bent, 1998). The mutant *chc2* was crossed with *DEX-CV-HA* line to obtain *chc2/DEX-CV-HA* line. *DEX-CV-HA* line was crossed with 3*5S::GAPC1-FLAG* transgenic line to obtain co-overexpression line *GAPC1-FLAG/DEX-CV-HA*. The triple knockdown lines (*amiR-CV/gapc1-1gapc2-1*) were obtained by crossing *amiR-CV-1* line with the double mutant *gapc1-1gapc2-1*. The homozygous triple knockdown lines were first selected by screening F2 generations with the combination of Basta and kanamycin, and then confirmed by the genotyping of the T-DNA insertion.

### Immunoprecipitation and LC-MS/MS-based protein identification

Seven-day-old seedlings of transgenic line *DEX-CV-HA* were cultured in MS/2 liquid medium containing 20µM DEX for 24h and then Concanamycin A (final concentration, 5µM) was added to the media for an additional 12h treatment. After treatment, the harvested seedlings were ground in liquid N_2_ and total proteins were extracted with a buffer [50mM Tris-Cl pH7.5, 150mM NaCl, 10%(v/v) glycerol, 1%(v/v) pvpp, 1%(v/v) Triton-X100, 1mM PMSF and protease inhibitor cocktail (Roche)] at 4°C for 3 h. The agarose beads conjugated with anti-HA monoclonal antibodies (Abmart, #M200135) were incubated with protein extractions for 2h at 4°C with a gentle shake. After washed for three times, the immunoprecipitated proteins were dissolved and denatured at 100°C and subjected to immunoblotting or LC-MS/MS analysis. LC-MS/MS analysis was performed as described previously (Shipman-Roston, Ruppel et al., 2010). Peptide identifications were accepted if the probability was >80% and protein identifications were accepted if there were three identical peptides with a >95% probability.

### Bimolecular Fluorescence Complementation (BiFC)

The DNA fragment of *c-myc-nYFP* (N-terminus of YFP with c-myc tag) was amplified from the vector pSPYNE-35S/pUC-SPYNE and fused to 3’ or 5’ terminus of *CV* and *CVΔC* to generate the fragments of *nYFP-CV*, *CV-nYFP*, and *CVΔC*, respectively. The *HA-cYFP* (C-terminus of YFP with HA tag) from vector pSPYCE-35S/pUC-SPYCE was fused to 5’ terminus of *CHC1*, *CHC2*, *GAPC1* and *GAPC2* or 3’ terminus of *SGR* to generate the fragment of *cYFP-CHC1*, *cYFP-CHC2*, *cYFP-GAPC1*, *cYFP-GAPC2,* and *SGR-cYFP*. All the fused fragments were ligated into the vector pEarly-Gate 100 (Earley, Haag et al., 2006). Vectors *pSPYNE-35S/pUC-SPYNE* and *pSPYCE-35S/pUC-SPYCE* were described previously (Walter, Chaban et al., 2004). For the transient expression, mesophyll protoplasts were isolated from four-week-old plants and were transformed by the above-mentioned BiFC constructs, according to a previous protocol (Yoo, Cho et al., 2007). BiFC fluorescence was excited at 488nm and detected at 525-552nm by using a confocal microscope (Leica TCS SP8). The primers are included in Table SII.

### Confocal microscopy observation of protein subcellular localization

Full-length cDNA of *CV* was amplified from cDNA of wild type mature leaves. The sequence of *GFP* was fused by PCR to 3’ or 5’ terminus of *CV*, respectively. The fragments of *CV-GFP* and *GFP-CV* were cloned into the XhoI-XbalI-digested vector pEarley-Gate 100 (Earley et al., 2006). By using the same strategy, DNA fragments of *mCherry-CHC2*, *GAPC1-mCherry*, *mCherry-DRP1A*, *OEP7-mCherry* were inserted into pEarley-Gate 100, respectively. The mesophyll protoplasts were transformed by the above-mentioned constructs as described previously (Yoo et al., 2007). A confocal laser scanning microscope (Leica TCS SP8) was used to perform fluorescence imaging. The excitation/emission wavelength for different fluorescence were: GFP (488nm/500 to 550nm), mCherry (543nm/585 to 615nm), chloroplast autofluorescence (405 nm/635 to 708nm). The primers are included in Table SII. Fluorescence colocalization was analyzed using PSC Colocalization plugin of ImageJ (French et al., 2008). Pearson’s and Spearman’s coefficient larger than 0.4 indicate colocalization, whereas lower or negative values indicate no colocalization (Belda-Palazon, Rodriguez et al., 2016).

### Co-IP

Three-week-old seedlings of transgenic lines *SSCU-GFP*, *DEX-CV-GFP*, *DEX-CVΔTM-GFP*, and *DEX-CVΔCD-GFP* were treated with 20µM DEX for 2 days. Total proteins were extracted and immunoprecipitated with agarose beads conjugated with anti-GFP antibodies (Abcam, #ab1240) as described above. Total proteins (Input) and co-IP samples were analyzed by immunoblotting with anti-GFP (ZENBIO China, #FG0407), anti-CHC (Agrisera, #AS10690-HRP) antibodies which can recognize both endogenous CHC1 and CHC2, and anti-GAPC antibodies (Agrisera, #AS152894), respectively.

The DNA fragment of *CV-HA* was inserted into pEarly-Gate 100. The fused fragments of *GAPC1-6×FLAG*, *GAPC2-6×FLAG*, and *6×FLAG-GFP* were inserted into the vector pBI121. These four constructs were introduced into *Agrobacterium tumefaciens* GV3101 strains for *Agrobacterium*-mediated transient expresssion in tobacco (*Nicotiana benthamiana*) leaves, as described previously (Li, Deng et al., 2016). The infiltrated tobacco leaves were harvested and extracted with extraction buffer after grinding in liquid N_2_. The agarose beads conjugated with anti-HA antibodies (Abmart, #M200135) were used for Co-IP analysis. Anti-CHC2, anti-GAPC, anti-GFP, anti-FLAG (Sigma, #A8592) and anti-HA (Abcam, #ab9110) were used for immunoblotting assays. The primers are included in Table SII.

### Immunoelectron microscopy

For the double immunolabeling analysis, 14-day-old *CPro::CV-GFP* plants were treated by 100mM mannitol and 3μM Concanamycin A for 12h. The leaves with expression of CV-GFP, which was confirmed by confocal microscope, were selected and fixed in paraformaldehyde (2%) and glutaraldehyde (2.5%). The ultrathin sections were immunolabeled with anti-GFP antibodies (from mouse) and anti-CHC antibodies (from rabbit), then treated with 5nm gold-conjugated goat anti-rabbit IgG and 20nm gold-conjugated goat anti-mouse IgG for 1h. The grids were stained with uranyl acetate and lead citrate and observed on a Phillips CM120 Biotwin. For the single immunolabeling analysis, the 2-week-old seedlings of *CPro::CV-GFP*, *CPro::CV-GFP/chc2*, and *CPro::CV-GFP/drp1a* were treated with 200mM mannitol for 24h. All the ultrathin sections from these lines were immunolabeled by anti-GFP antibodies and goat anti-mouse secondary antibodies conjugated 10nm gold particles.

### Isolation of chloroplasts and envelope membranes

Three-week-old *DEX-CV-GFP* seedlings were treated with 10μM DEX for 2d. The chloroplasts were isolated from leaves by using Minute^TM^ chloroplast isolation kit (Invent Biotechnologies, Inc). The inner and outer envelope fractions from the isolated chloroplasts were separated as described previously (Wang et al., 2014). Total proteins, isolated chloroplasts, outer and inner envelop membranes were dissolved in 6×loading buffer and separated by SDS-PAGE for immunoblotting analysis. The antibodies raised against TIC110 (PhytoAB Inc, PHY0449S), TOC33 (PhytoAB Inc, PHY0143), PsbO (Agrisera, PHY0094A), and GS1/GS2 (Agrisera, AS08295) were chosen to detect the representative proteins in all the fractions.

### RNA Extraction and Quantitative RT-PCR

TRNzol^+R^ Reagent (Life Technologies Corporation) was used to isolate high-quality total RNAs from the different seedlings. Total RNAs were digested with gDNA Eraser to eliminate genomic DNA. One μg of total RNAs were used for reverse-transcription reactions, according to the manufacturer’s protocol of PrimeScriptTM RT reagent Kit with gDNA Eraser (Takara, #RR047A). The qPCR assay was performed by using UltraSYBR Mixture (KWBIO, #CW2602) on the CFX Connect^TM^ Real-Time PCR Detection System (Bio-Rad). *ACTIN* was used as a normalization control and the relative expression of each test gene was calculated by the 2^-ΔΔCT^ method (Livak & Schmittgen, 2001). The primers of qRT-PCR are included in Table SII.

### Water stress experiments

For water deficit stress assays, 7-day-old Arabidopsis seedlings were transplanted into pots and grown for 7d. Watering was withheld for 12d (to *chc2/DEX-CV-HA*, *DEX-CV-HA*, Col-0, *amiR-CV-1*, *gapc1gapc2*, and *amiR-CV-1/gapc1gapc2* plants) or 14d (to Col-0, *chc2*, *DEX-CV-HA*, *GAPC1-FLAG/DEX-CV-HA* plants). Survival rate and chlorophyll contents was analyzed 3d after re-watering.

## Supplementary data

Supplemental Information includes six figures and two tables, and can be found with this article online.

## Acknowledgements

This work was supported by National Natural Science Foundation of China (Grant No. 31871354), the Hundred Talents Program (Chinese Academy of Sciences), Startup Funding of Anhui Agricultural University, and the “Wanjiang Scholar” Program (Anhui Province).

## Author contributions

S. W. and E. B. designed the experiments. T. P., Y. L., C. L., Y. T., W. T., L. G., C. K., J. F., and H. L. conducted the experiments. T. P. and S. W. analyzed the data. S. W., E. B., Y. L., and T. P. wrote the manuscript.

## Supplemental Figures and Tables

**Figure S1:**
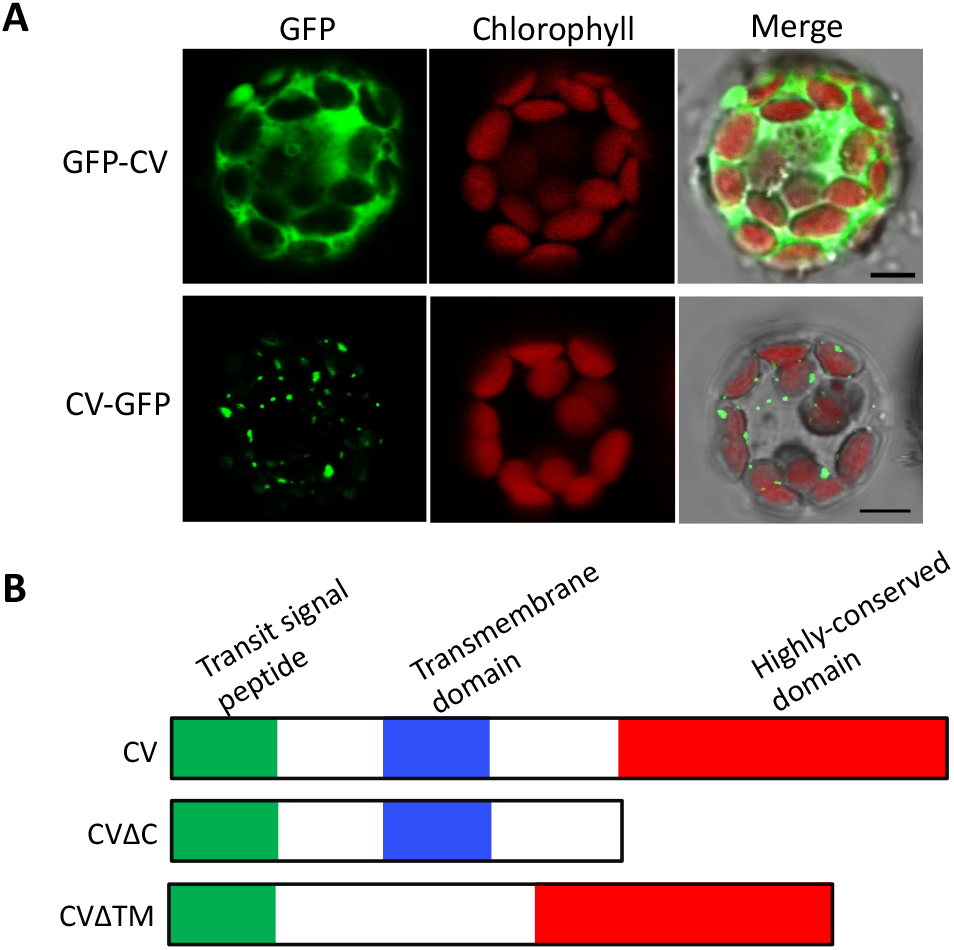
GFP-CV shows no dot-like structure. (A) Confocal microscopic observation of GFP-CV and CV-GFP transiently expressed in the protoplasts isolated from wild type leaves (Col-0). Bar=5μm (B) The protein domains of CV. The green is the transient signal peptide. The blue is the transmembrane domain. The red is a previously-unknown domain which is highly conserved in the plant kingdom.

**Figure S2:**
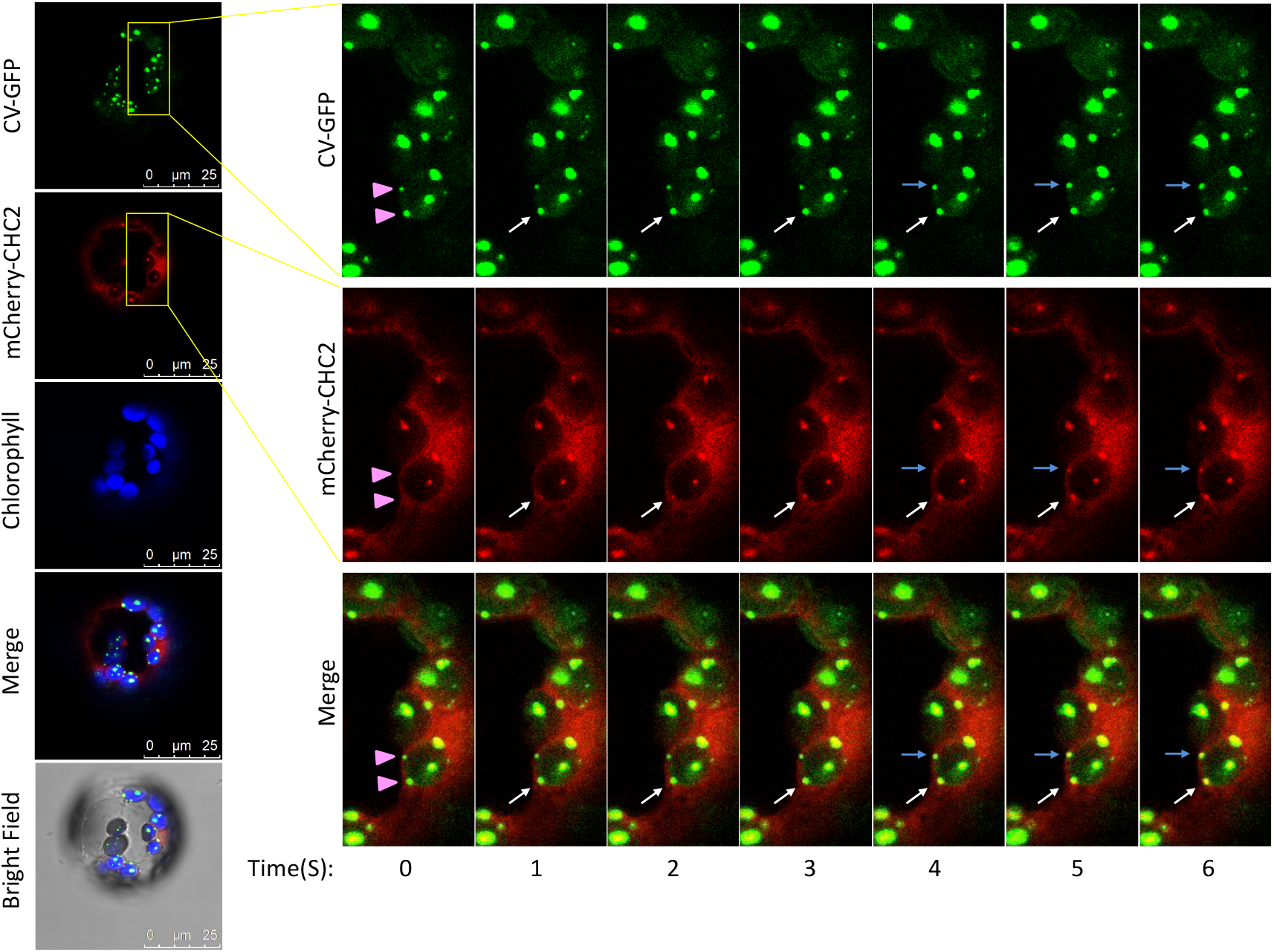
Time-lapse observation that CV interacts CHC2. *CV-GFP* and *mCherry-CHC2* were transiently expressed in the protoplasts isolated from Col-0 plants. The right time-lapse observations were amplified from the yellow frames in the left pictures. Purple triangles indicated that the CV-GFP-labeled “dots” were not associated with mCherry-CHC2 in the first place. The white and blue arrows indicated the moments that mCherry-CHC2 was recruited.

**Figure S3:**
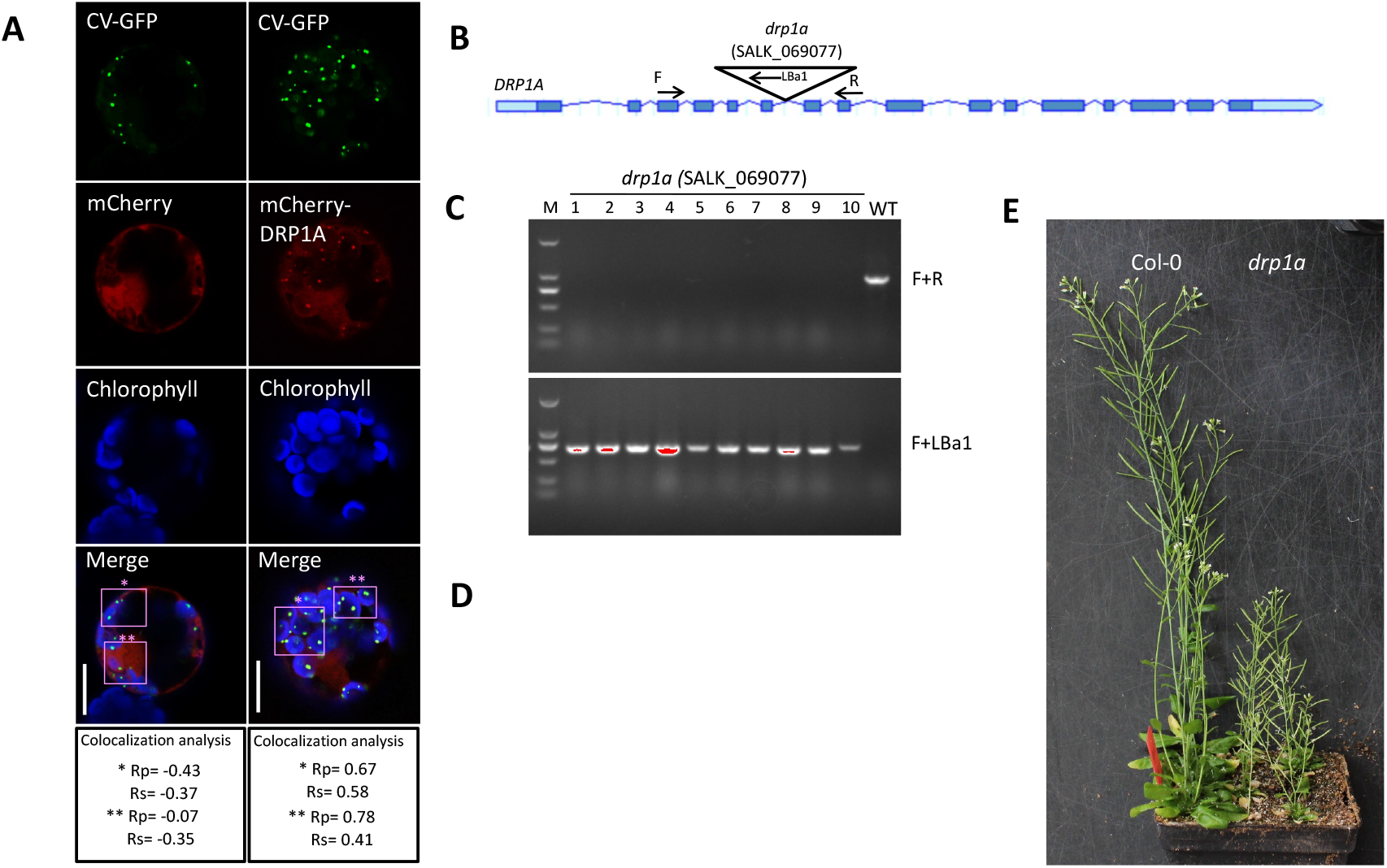
Co-localization of CV and DRP1A and Identification of T-DNA insertion mutant *drp1a*. (**A**) The protoplasts isolated from leaves of wild type plants were co-transformed by the vector *CV-GFP* with mCherry-DRP1A or mCherry. Bar=10μm. The bottom panels showed the colocalization analysis for the purple boxed region. Pearson’s (Rp) and Spearman’s (Rs) coefficients of “green” and “red” channels were calculated (n>10). The Rp and Rs values in the range +0.4 to 1 indicate colocalization, whereas low or negative values indicate lack of colocalization. (**B**) T-DNA of the mutant *drp1a* (germplasm SALK_069077), was inserted in the sixth intron of *DRP1A* gene. (**C**) PCR verification of T-DNA insertion in the genome of *drp1a*. (**D**) The qRT-PCR analysis of *DRP1A* gene expression in the seedlings of Col-0 and *drp1a*.

**Figure S4:**
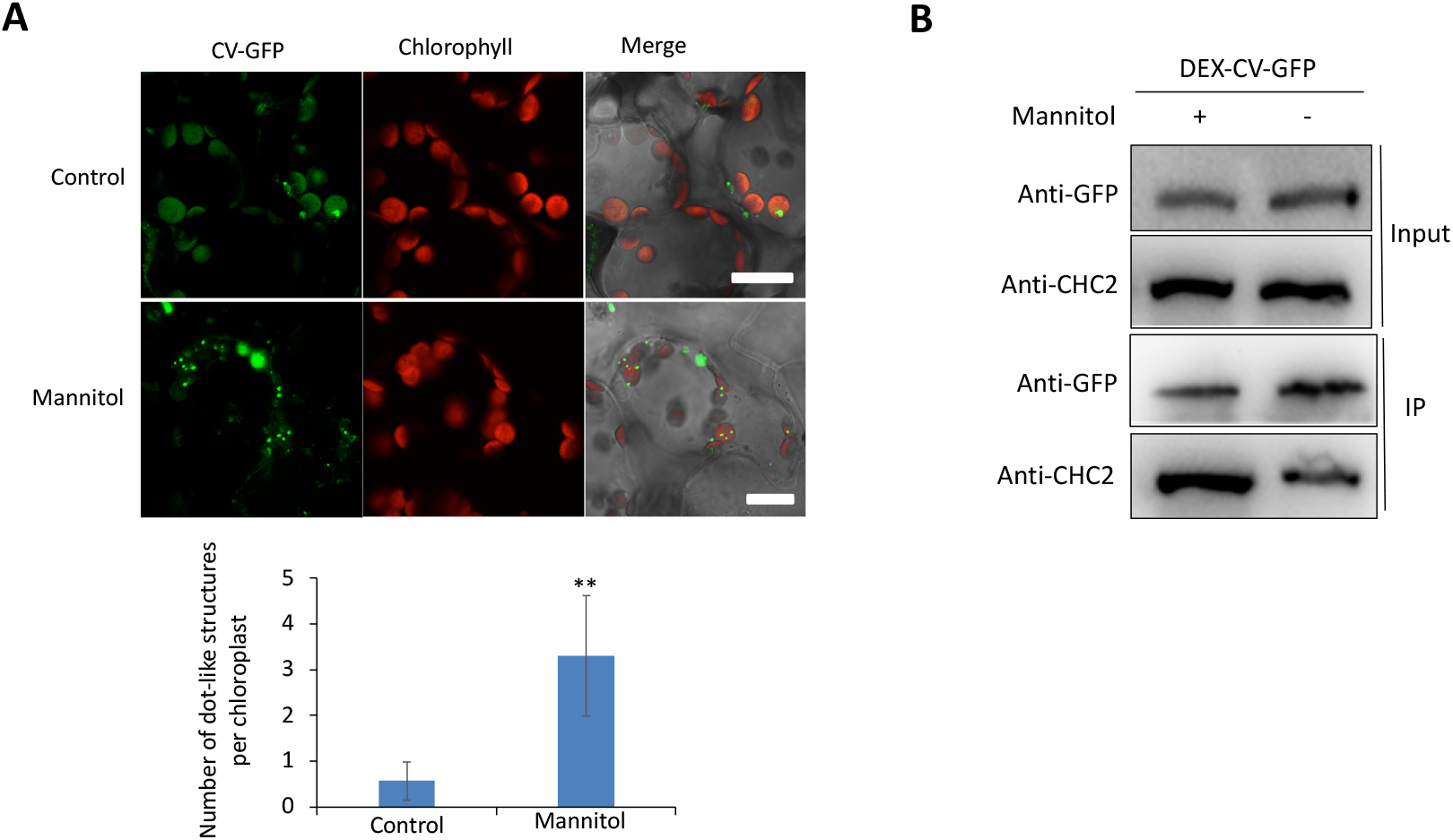
Osmotic stress enhanced CVV formation and the CV-CHC2 interaction. (**A**) The transgenic DEX-CV-GFP plants were treated with 10μM DEX in combination with 200mM mannitol or without (control) for 12h. The mesophyll cells were observed by confocal microscope. Bar=20μm. Means±SD of the number of CV-GFP dot-like structures was obtained from more than 50 chloroplasts in three independent experiments. Asterisks “**” indicate significant difference at P<0.001. (**B**) The DEX-CV-GFP plants after the treatment as described in (**A**) were used for Co-IP assays with beads conjugated with anti-GFP antibodies. The Input and IP samples were detected by the immunoblotting with anti-GFP and anti-CHC antibodies, respectively.

**Figure S5:**
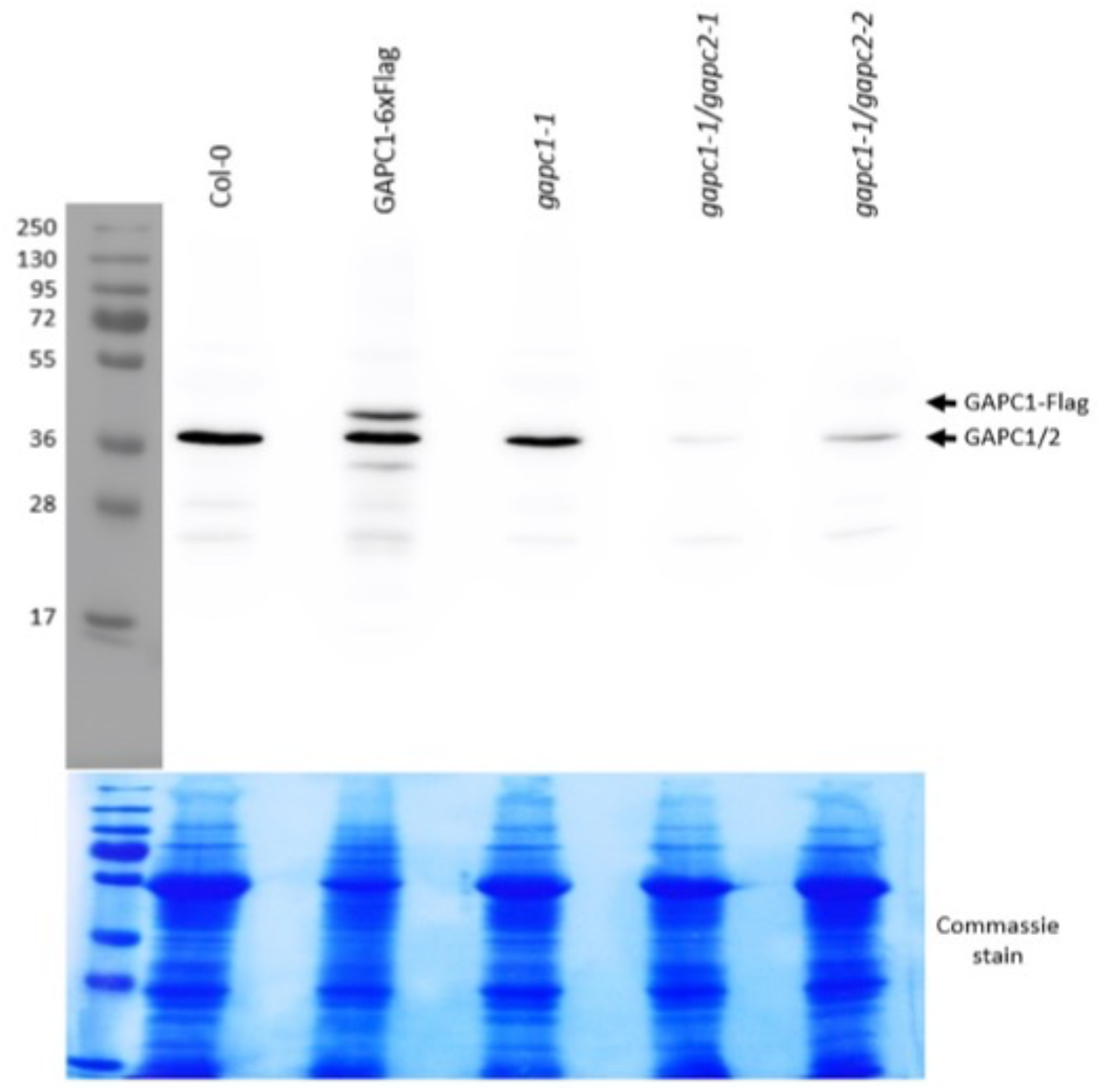
Immunoblot analysis of single and double *gapc* mutants. Total proteins extracted from 3-week-old seedlings of Col-0, GAPC1-FLAG, *gapc1-1*, *gapc1-1/gapc2-1*, and *gapc1-1/gapc2-1* were immunoblotted with anti-GAPC antibodies which can recognize both GAPC1 and GAPC2.

**Figure S6:**
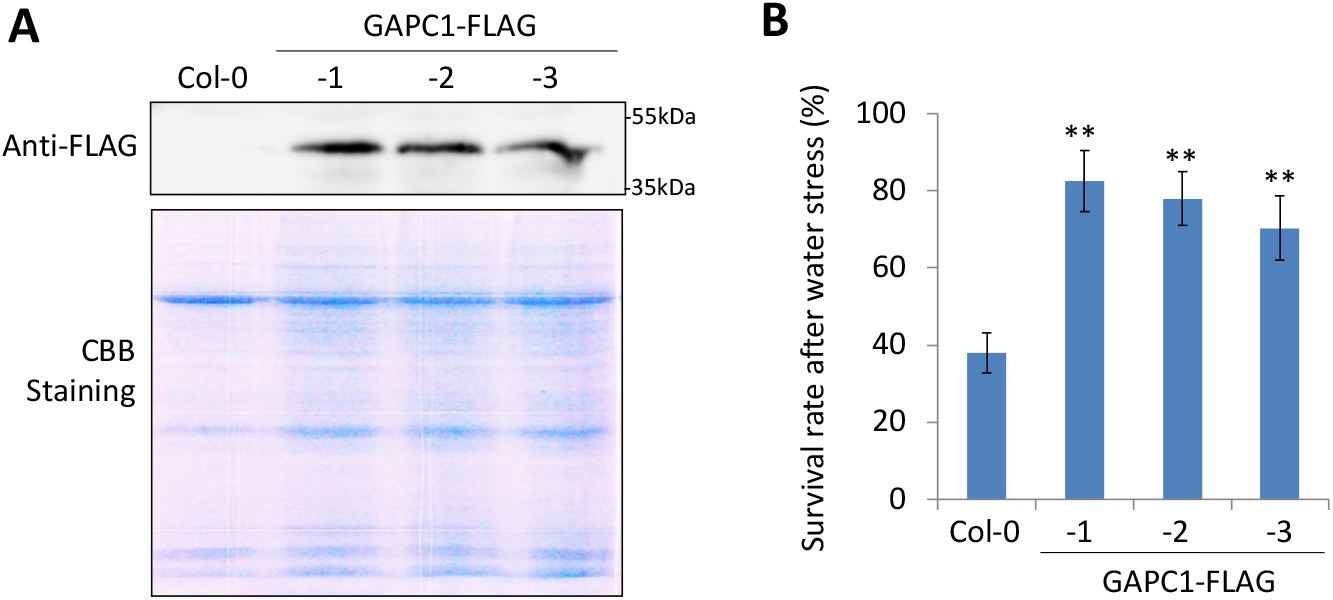
The GAPC1-FLAG overexpression lines showed drought tolerance. (**A**) Immunoblotting analysis of Col-0 and three *GAPC1-FLAG* overexpression lines with anti-FLAG antibodies. (**B**) Two-week-old seedlings of Col-0 and three *GAPC1-FLAG* overexpression lines were subjected to water stress for 14 d followed by rewatering for 3 d. Survival rate was determined 3 d after rewatering. Eighteen plants for each line were evaluated and Mean ± SD were obtained from three independent tests. Asterisks “**” indicate significant difference at P<0.001 from Col-0.

**Table SI:** Potential interactors of CV as identified by Co-IP and LC-MS/MS.

| Uniport | Protein names | Gene names | MW [kDa] | Protein score | Sequence coverage (%) | # Peptides |
| --- | --- | --- | --- | --- | --- | --- |
| Q9ASS6 | Peptidyl-prolyl cis-trans isomerase CYP20-2 | CYP20-2 | 28 | 749.85 | 32.37 | 24 |
| Q8S8K8 | CHLOROPLAST VESICULATION | CV | 16 | 654.1 | 37.21 | 17 |
| P23321 | Oxygen-evolving enhancer protein 1-1 | PSBO1 | 35 | 390.17 | 22.88 | 16 |
| O22609 | Protease Do-like 1 | DEGP1 | 47 | 318.46 | 31.41 | 14 |
| P25858 | Glyceraldehyde-3-phosphate dehydrogenase | GAPC | 37 | 267.25 | 27.49 | 14 |
| O50008 | 5-methyltetrahydropteroyltriglutamate-homocysteine methyltransferase | CIMS | 84 | 204.94 | 10.94 | 13 |
| Q9LTX9 | Heat shock 70 kDa protein 7 | HSP70-7 | 77 | 186.23 | 14.89 | 12 |
| F4JBY2 | Transketolase | At3g60750 | 80 | 186.84 | 12.01 | 11 |
| Q96300 | 14-3-3-like protein GF14 | GRF7 | 30 | 147.82 | 29.81 | 11 |
| Q9LD57 | Phosphoglycerate kinase 1 | PGK1 | 50 | 142.47 | 19.05 | 11 |
| Q9LVM2 | Oxidoreductase | NDHN | 23 | 112.47 | 16.3 | 10 |
| P43297 | Cysteine proteinase RD21a | RD21A | 51 | 111.81 | 6.69 | 10 |
| Q0WLB5 | Clathrin heavy chain 2 | CHC2 | 192 | 88.72 | 2.03 | 10 |
| P25857 | Glyceraldehyde-3-phosphate dehydrogenase B | GAPB | 48 | 86.36 | 11.36 | 9 |
| Q42547 | Catalase-3 | CAT3 | 57 | 85.34 | 9.47 | 9 |
| P53492 | Actin-7 | ACT7 | 42 | 84.2 | 10.21 | 8 |
| Q9XF88 | Chlorophyll a-b binding protein CP29.2 | LHCB4.2 | 31 | 82.01 | 15.23 | 8 |
| Q56YW9 | Tubulin beta-2 chain | TUB2 | 51 | 81.45 | 11.83 | 7 |
| Q9LJG3 | GDSL esterase/lipase | ESM1 | 44 | 80.32 | 12.85 | 7 |
| O23654 | V-type proton ATPase catalytic subunit A | VHA-A | 69 | 79.48 | 10.02 | 7 |
| Q39249 | Violaxanthin de-epoxidase | VDE1 | 52 | 78.97 | 26.21 | 6 |
| Q9SYI0 | Protein translocase subunit SECA1 | SECA1 | 115 | 78.68 | 4.03 | 5 |
| Q9C8G5 | Early-responsive to dehydration stress protein | ERD4 | 82 | 69.41 | 12.67 | 5 |
| P11574 | V-type proton ATPase subunit B1 | VHA-B1 | 54 | 66.74 | 5.88 | 5 |
| Q9SHE8 | Photosystem I reaction center subunit III | PSAF | 24 | 64.38 | 15.07 | 5 |
| B9DGT7 | Tubulin alpha-2 chain | TUA2 | 50 | 62.54 | 16.05 | 5 |
| Q9LI74 | Protein CHUP1 | CHUP1 | 112 | 61.26 | 14.53 | 4 |
| Q9SMX3 | Mitochondrial outer membrane protein porin 3 | VDAC3 | 29 | 59.57 | 2.05 | 4 |
| Q9S841 | Oxygen-evolving enhancer protein 1-2 | PSBO2 | 35 | 58.98 | 3.25 | 4 |
| P21240 | Chaperonin 60 subunit beta 1 | CPNB1 | 64 | 58.69 | 2.13 | 4 |
| F4JV51 | ENTH/VHS/GAT family protein | At4g32760 | 73 | 58.16 | 3.75 | 4 |
| P23586 | Sugar transport protein 1 | STP1 | 58 | 57.19 | 4.28 | 4 |
| O03986 | Heat shock protein 90-4 | HSP904 | 80 | 57.12 | 2.86 | 4 |
| Q9SA52 | Chloroplast stem-loop binding protein | CP41B | 41 | 51.15 | 5.85 | 4 |
| Q8LPR9 | Protein TIC110 | TIC110 | 112 | 50.98 | 10.23 | 4 |
| Q9M0E5 | Vacuolar protein Sorting 15 | VPS15 | 168 | 50.23 | 1.03 | 3 |
| Q01525 | 14-3-3-like protein GF14 omega | GRF2 | 29 | 49.88 | 18.18 | 3 |
| Q39258 | V-type proton ATPase subunit E1 | VHA-E1 | 26 | 48.84 | 12.14 | 3 |
| Q8LCE1 | Glutamine synthetase cytosolic isozyme 1-2 | GS2 | 39 | 47.97 | 14.38 | 3 |
| F4HW17 | Endomembrane protein 70-like protein | At1g08350 | 68 | 47.61 | 3.42 | 3 |
| O03042 | Ribulose biphosphate carboxylase large chain | rbcL | 53 | 429.28 | 48.85 | 9 |
| P04778 | Chlorophyll a-b binding protein 1, chloroplastic | LHCB1.3 | 28 | 671.49 | 48.31 | 7 |
| Q9S7M0 | Chlorophyll a-b binding protein 3, chloroplastic | LHCB3 | 29 | 383.14 | 22.26 | 3 |
| Q9LMQ2 | Chlorophyll a-b binding protein, chloroplastic | F7H2.16 | 28 | 280.03 | 22.48 | 4 |
| P36212 | 50S ribosomal protein L12-3, chloroplastic | RPL12C | 20 | 86.43 | 7.49 | 1 |
| Q9LW57 | Probable plastid-lipid-associated protein 6, chloroplastic | PAP6 | 30 | 58.49 | 7.75 | 2 |
| P49201 | 40S ribosomal protein S23-2 | RPS23B | 16 | 51.21 | 7.75 | 1 |
| P93819 | Malate dehydrogenase 1, cytoplasmic | MDH1 | 36 | 48.84 | 3.61 | 1 |
| P93285 | Cytochrome c oxidase subunit 2 | COX2 | 30 | 47.78 | 7.69 | 1 |
| Q9LR64 | Photosystem II repair protein PSB27-H1, chloroplastic | PSB27-1 | 19 | 44.06 | 5.75 | 1 |

**Table S2.**
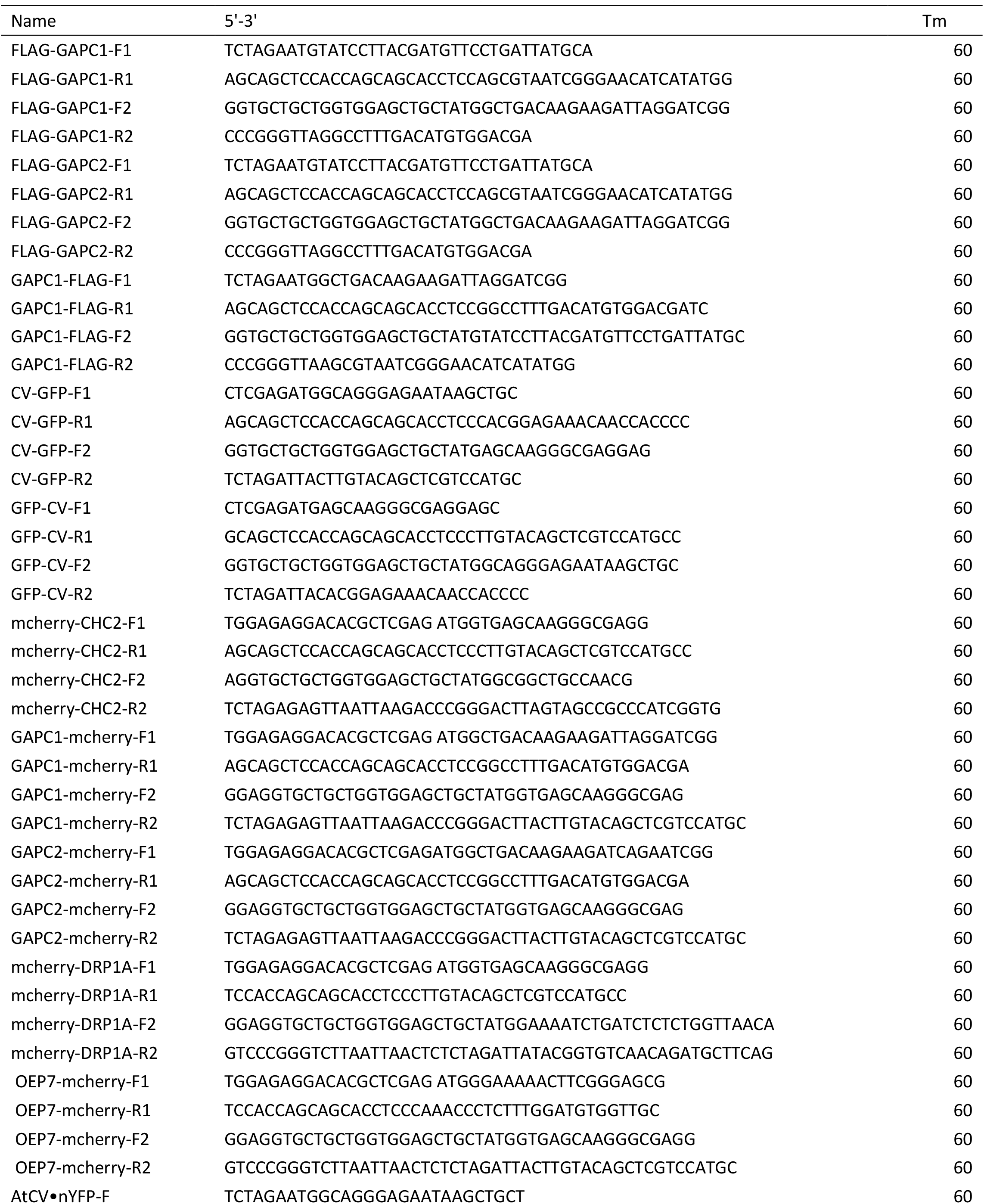
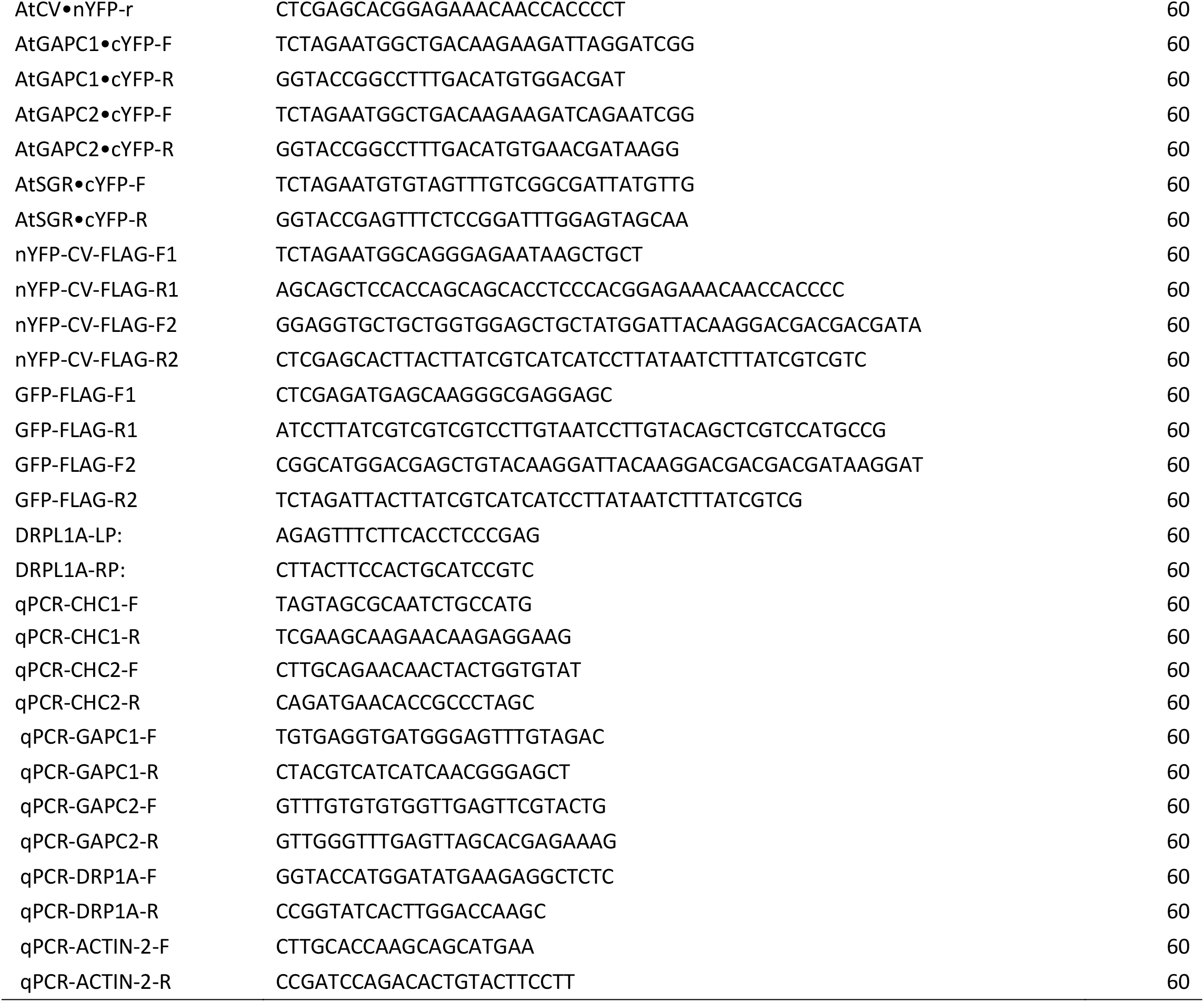
Sequence of primers used in this study.

